# An armored marine reptile from the Early Triassic of South China

**DOI:** 10.1101/2022.10.17.512493

**Authors:** Andrzej S. Wolniewicz, Yuefeng Shen, Qiang Li, Yuanyuan Sun, Yu Qiao, Yajie Chen, Jun Liu

## Abstract

Sauropterygia was a taxonomically and ecomorphologically diverse clade of Mesozoic marine reptiles spanning the Early Triassic to the Late Cretaceous. Sauropterygians are traditionally divided into two groups representing two markedly different body plans – the short-necked, durophagous Placodontia and the long-necked Eosauropterygia – whereas Saurosphargidae, a small clade of marine reptiles possessing a dorsal ‘rib-basket’, is considered as the sauropterygian sister-group. However, the early evolutionary history of sauropterygians and their phylogenetic relationships with other groups within Diapsida are still incompletely understood. Here, we report a new saurosphargid from the Early Triassic of South China – *Prosaurosphargis yingzishanensis* gen. et sp. nov. – representing the earliest known occurrence of the clade. An updated phylogenetic analysis focussing on the interrelationships within diapsid reptiles recovers saurosphargids as nested within sauropterygians, forming a clade with eosauropterygians to the exclusion of placodonts. Furthermore, a clade comprising *Eusaurosphargis* and *Palatodonta* is recovered as the sauropterygian sister-group within Sauropterygomorpha tax. nov. The phylogenetic position of several Early and Middle Triassic sauropterygians of previously uncertain phylogenetic affinity, such as *Atopodentatus, Hanosaurus, Majiashanosaurus* and *Corosaurus*, is also clarified, elucidating the early evolutionary assembly of the sauropterygian body plan. Finally, our phylogenetic analysis recovers Testudinata and Archosauromorpha within Archelosauria, a result strongly supported by molecular data, but until now not recovered by any phylogenetic analysis using a morphology-only data set. Our study provides evidence for the rapid diversification of sauropterygians in the aftermath of the Permo-Triassic mass extinction event and emphasises the importance of broad taxonomic sampling in reconstructing phylogenetic relationships among extinct taxa.

## Introduction

Several groups of reptiles invaded the marine realm in the aftermath of the Permo-Triassic mass extinction (PTME), the largest extinction event in Earth’s history (Benton 2015). This phenomenon was likely a result of the scarcity of marine competitors and predators caused by the PTME and high productivity in the incipient shallow marine environment (Vermeij and Motani 2018). Triassic marine reptiles, including the iconic Ichthyosauromorpha and Sauropterygia, as well as some other smaller and lesser known groups, achieved high taxonomic and ecological diversity rapidly after their emergence in the late Early Triassic and played a pivotal role in the reorganisation of marine food webs following the PTME (Scheyer et al. 2014; Kelley and Pyenson 2015; Motani et al. 2015, 2017; Jiang et al. 2016; Stubbs and Benton 2016; Cheng et al. 2019; Moon and Stubbs 2020; Li and Liu 2020; Sander et al. 2021). Because Mesozoic marine reptiles represent likely several independent transitions from a terrestrial to an aquatic lifestyle, they also provide ideal systems to analyse the roles of function and constraint in determining evolutionary pathways (Motani 2009; Benson 2013).

Sauropterygia was a diverse clade of Mesozoic marine reptiles that first appeared in the late Early Triassic (Jiang et al. 2014; Li and Liu 2020) and remained important predators in marine ecosystems until their extinction at the end of the Late Cretaceous (Bardet 1994; Rieppel 2000a; Benson and Druckenmiller 2014). Sauropterygia is traditionally divided into two major lineages, representing two markedly different body plans – the Placodontia and the Eosauropterygia, the latter comprising Pachypleurosauria, Nothosauroidae and Pistosauroidea (Rieppel 2000a). Placodonts were characterised by the presence of short necks and short skulls and possessed crushing palatal dentition almost certainly used for feeding on hard-shelled invertebrates (Rieppel 2002; Crofts et al. 2017). Some early-diverging placodonts had a limited covering of osteoderms on their backs and possibly limbs (Jiang et al. 2008; Klein and Scheyer 2013), but derived forms evolved extensive dorsal armor superficially similar to that of turtles (Scheyer 2007; Wang et al. 2020). Eosauropterygians, on the other hand, had elongated necks and elongated skulls with pointed dentition suitable for capturing fast-moving prey (Rieppel 2000a, 2002). Placodonts remained restricted to shallow marine environments until their extinction in the Late Triassic, whereas eosauropterygians evolved a suite of adaptations for a pelagic lifestyle and became one of the dominant groups of marine reptiles in the Jurassic and Cretaceous (Kelley et al. 2014).

Even though sauropterygians have a rich fossil record and a long history of scientific research, the early evolution of the group is still incompletely understood. In a broad phylogenetic context, sauropterygians have been consistently recovered within Diapsida, but their exact phylogenetic position relative to other diapsid groups remains unresolved. Several diapsid clades, including Saurosphargidae (Chen et al. 2014; Li et al. 2014, 2018; Neenan et al. 2015; Wang et al. 2022), Testudinata (de Braga and Rieppel 1997; Schoch and Sues 2015), Ichthyosauromorpha (Martínez et al. 2021) and the enigmatic reptile *Eusaurosphargis dalsassoi* (Scheyer et al. 2017) were proposed as the sauropterygian sister-group, but its exact identity is still a matter of debate. Because sauropterygians were proposed as being closely related not only to turtles, but also to archosauromorphs (Chen et al. 2014; Neenan et al. 2015; Martínez et al. 2021), resolving their phylogenetic placement within diapsids is of crucial importance for solving the phylogenetic uncertainty surrounding Archelosauria – a clade comprising turtles and archosauromorphs strongly supported by molecular data but thus far not recovered by any phylogenetic analysis using a morphology-only dataset (Crawford et al. 2015; Lyson and Bever 2020).

Saurosphargids are a small clade of Mesozoic marine reptiles characterised by the presence of a dorsal ‘rib-basket’ and a moderately- to well-developed osteoderm covering (Li et al. 2011, 2014). Saurosphargids are known from the Middle Triassic of Europe and South China, although recent evidence suggests they could have survived as late as the Late Triassic (Scheyer et al. 2022). Saurosphargids comprise as many as four taxa – *Saurosphargis volzi* (considered a nomen dubium by some authors) (Huene 1936; Nosotti and Rieppel 2003; Scheyer et al. 2017), the heavily armored *Sinosaurosphargis yunguiensis* (Li et al. 2011; Hirasawa et al. 2013) and two species in the genus *Largocephalosaurus* (*L. polycarpon* and *L. qianensis*) (Cheng et al. 2012; Li et al. 2014). Saurosphargids are one of the reptile groups proposed as the sister-group of sauropterygians (see above), but some recent phylogenetic analyses have suggested their placement within sauropterygians instead, as either the sister-group to placodonts (Schoch and Sues 2015; Simões et al. 2018; Martínez et al. 2021) or eosauropterygians (Scheyer et al. 2017; Wang et al. 2019).

Our understanding of the early evolution of the sauropterygian body plan is also hindered by the uncertain phylogenetic position of several Early and Middle Triassic taxa. *Hanosaurus hupehensis* (Young 1972; Rieppel 1998a; Wang et al. 2022) and *Majiashanosaurus discocoracoidis* (Jiang et al. 2014) from the Early Triassic of South China are variably recovered as either outside of the clade comprising Saurosphargidae + Sauropterygia (*Hanosaurus*; Wang et al. 2022), as pachypleurosaurs (Jiang et al. 2014; Neenan et al. 2015; Lin et al. 2021; Wang et al. 2022) or as outgroups to a clade comprising nothosauroids and pachypleurosaurs to the exclusion of pistosauroids (Li and Liu 2020). The phylogenetic placement of *Corosaurus alcovensis* from the Early–Middle Triassic of Wyoming, USA, is also unresolved, with difrerent authors arguing for its early-diverging eosauropterygian (Rieppel 1994; Li and Liu 2020), eusauropterygian (more closely related to nothosauroids and pistosauroids than to pachypleurosaurs) (Neenan et al. 2015; Lin et al. 2021) or pistosauroid (Rieppel 1998b) affinity. The phylogenetic position of the herbivorous hammer-headed sauropterygian *Atopodentatus unicus* from the Middle Triassic of South China relative to placodonts and eosauropterygians also remains uncertain (Cheng et al. 2014; Li et al. 2016). These conflicting phylogenetic placements are likely the result of inadequate sampling of Early and Middle Triassic sauropterygian ingroup taxa, as well as the inclusion of only a limited number of diapsid reptile clades, including potential sauropterygian outgroups, in phylogenetic analyses.

Here, we report an Early Triassic saurosphargid from South China, representing the earliest recorded occurence of the group. We also present an updated phylogenetic hypothesis for Diapsida with a particular focus on Sauropterygia, which we use as context for discussing the early evolutionary assembly of the sauropterygian body plan.

## Results

### Geological background

Yingzishan quarry, where the new specimen was collected from, is located on the northern boundary of the Yangtze Platform (Figure 1A) (Li and Liu 2020). The new specimen represents the Early Triassic Nanzhang-Yuan’an marine reptile fauna (Li and Liu 2020) and is preserved in dark grey laminated and thin-bedded carbonate mudstone with some carbonaceous interactions from the upper part of the third member of the Jialingjiang Formation (Figure 1B). There are also some peloids, replacive dolomites and microbial mats in the fossiliferous levels (Chen et al. 2022). Based on the published sedimentological accounts (Wang et al. 2011; Chen et al. 2016b; Yan et al. 2018, 2021; Li et al. 2020; Zhao et al. 2022) and field observations, a restricted, stagnant, and hypersaline lagoon within a tidal flat environment is inferred as the burial setting of the marine reptiles of the Nanzhang-Yuan’an fauna (Chen et al. 2022).

**Figure 1.**
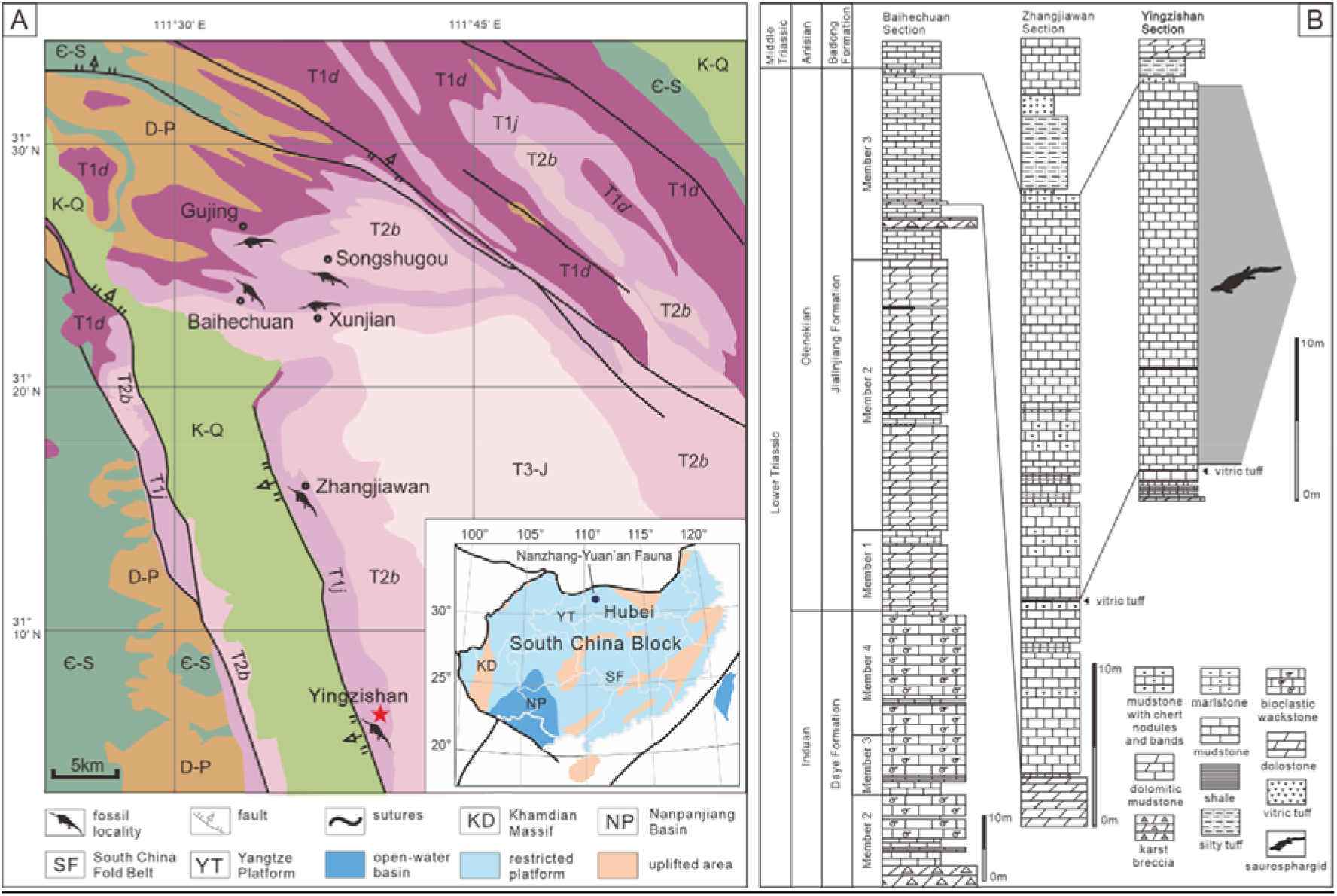
Locality and horizon of *Prosaurosphargis yingzishanensis* (HFUT YZSB-19-109). (A)The geological map of the Nanzhang-Yuan’an region showing Yingzishan quarry where HFUT YZSB-19-109 was collected; inset is a paleogeographic map of the South China Block in the Triassic showing the location of the Nanzhang-Yuan’an fauna (after Qiao et al. 2020). (B) Stratigraphic column showing the horizon from which HFUT YZSB-19-109 was collected. Abbreviations: Є-S, Cambrian–Silurian; D-P, Devonian–Permian; K-Q, Cretaceous–Quaternary; T1d, Daye Formation, Lower Triassic; T1j, Jialingjiang Formation, Lower Triassic; T2b, Badong Formation, Middle Triassic; T3-J, Upper Triassic–Jurassic.

### Systematic paleontology

Archelosauria Crawford et al. 2015

Sauropterygomorpha tax. nov.

urn:lsid:zoobank.org:pub:[…] (taxon will be registered in ZooBank after paper acceptance)

#### Definition

The most recent common ancestor of *Eusaurosphargis dalsassoi* and *Pistosaurus longaevus* and all of its descendants.

#### Diagnosis

Osteoderms present (ch. 143.1), body strongly flattenned dorso-ventrally (ch. 144.1), clavicle applied to the medial surface of scapula (ch. 148.1), metatarsal V long and slender (ch. 186.0), metatarsal I less than 50% the length of metatarsal IV (ch. 189.1).

Sauropterygia Owen, 1860

Saurosphargidae Li et al., 2011

*Prosaurosphargis yingzishanensis* gen. et sp. nov.

urn:lsid:zoobank.org:pub:[…] (taxon will be registered in ZooBank after paper acceptance)

#### Holotype

HFUT YZSB-19-109, a partial postcranial skeleton (Figure 2). The specimen is housed in the Geological Museum of Hefei University of Technology, Hefei, Anhui Province, China (HFUT).

**Figure 2.**
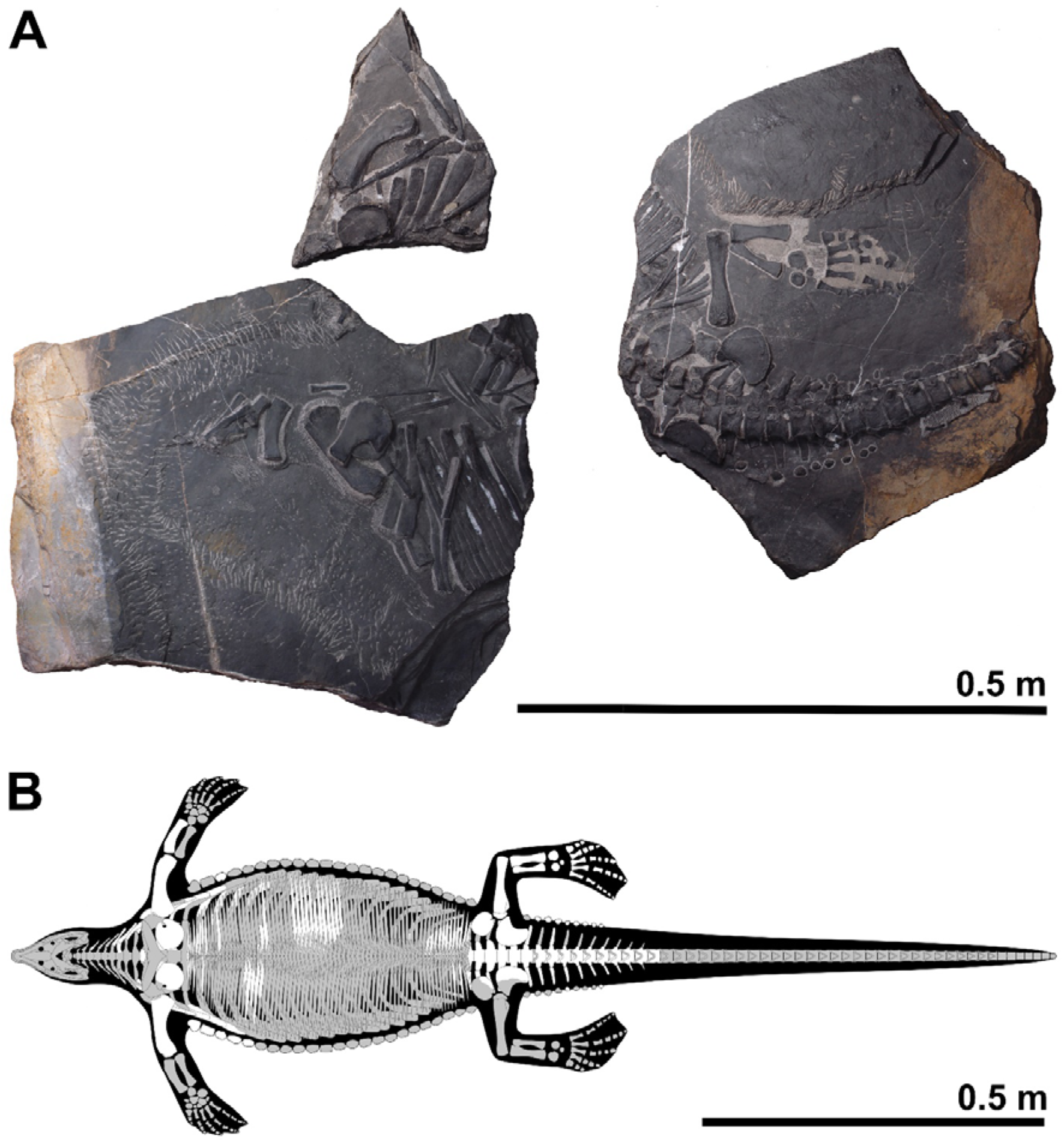
Holotype of *Prosaurosphargis yingzishanensis* (HFUT YZSB-19-109) (A) and skeletal reconstruction with known elements highlighted in white (B).

#### Etymology

Genus name from the Greek preposition πρó (pró) meaning before and *Saurosphargis*, the name of the type genus of the family Saurosphargidae (Li et al. 2011). The specific epithet refers to the type locality (see above).

#### Horizon and locality

Third Member of the Jialingjiang Formation (uppermost Spathian, Olenekian, Lower Triassic), Yingzishan quarry, Yuan’an County, Hubei Province, China.

Stratigraphic note: Traditionally, the Jialingjiang Formation has been divided into four members by most authors studying the region (Li et al. 2002; Zhao et al. 2008; Cheng et al. 2015). From base to top, the first member consists of thick-bedded to massive dolostone, the second member consists of vermicular limestone intercalated with dolostone, the third member consists of laminated thin- to medium-bedded limestone to dolostone, while the fourth member is composed of dolostone and karstified breccia. Thus, in the four-member division of the Jialingjiang Formation, the thick volcanic ash (Figure 1B) marks the bottom of the fourth member. However, recent geological mapping in the region (Chen et al. 2016a, b) proposed that the fourth member of the Jialingjiang Formation should be included in the Middle Triassic Badong Formation, indicating a three-member division of the Jialingjiang Formation. This division is consistent with the lithology of the Jianlingjiang and Badong formations as defined by the official geological guide of Hubei Province (Bureau of Geology and Mineral Resources of Hubei Province 1990). The division of the Lower and Middle Triassic in the region is also consistent with the widespread thick volcanic ash as a marker of the Lower–Middle Triassic boundary in the Yangtze Platform (Chen et al. 2020). Consequently, this division was followed by Cheng et al. (2019), Li and Liu (2020), and Qiao et al. (2020). Nevertheless, recently the same group of authors (Yan et al. 2021, contra. Chen et al. 2016a, b and Cheng et al. 2019) redefined the second and third members of the Jialingjiang Formation in the Nanzhang-Yuan’an region. They regarded the lowermost base of the thick vitric tuff as the boundary of the second and third members. Thus, the typical Nanzhang-Yuan’an fauna was placed into the newly defined second member of the Jialingjiang Formation (Cheng et al. 2022; Zhao et al. 2022). We argue that this new definition and division of different members of the Jialingjiang Formation contradicts the official geological guide (Bureau of Geology and Mineral Resources of Hubei Province 1990) and causes confusion. Therefore, we prefer to maintain the definition of the thick volcanic ash as the boundary between the Early Triassic Jialingjiang and the Middle Triassic Badong formations in the Nanzhang-Yuan’an region (Figure 1B), pending future updates of the official geological guide of Hubei Province.

#### Diagnosis

A saurosphargid characterised by the following combination of character states: 1) spaces between dorsal transverse processes anteroposteriorly shorter than the anteroposterior widths of the transverse processes (like in *Saurosphargis* and *Sinosaurosphargis*, but different from *Largocephalosaurus*, in which the spaces between the dorsal transverse processes are wider than the transverse processes); 2) ribs without uncinate processes (like in *Sinosaurosphargis*, but unlike *Saurosphargis* and *Largocephalosaurus*, in which uncinate processes are present); 3) three sacral vertebrae/ribs (like in *L. polycarpon*, compared with two sacrals present in *L. qianensis*) (number of sacral vertebrae unknown in *Sinosaurosphargis*); 4) osteoderms forming a median and lateral rows (and possibly also parasaggital rows) (similar to *L. polycarpon*, but different from *L. qianensis*, in which additional, small osteoderms more extensively cover the lateral sides of the body and different from *Sinosaurosphargis*, in which the osteoderms form an extensive dorsal armor); 5) ectepicondylar groove on humerus present (like in *L. qianensis*, but unlike *L. polycarpon*); 6) entepicondylar foramen in humerus absent (as in *L. qianensis*, different from *L. polycarpon*) (details of humerus morphology unknown in *Sinosaurosphargis*); 7) radius short relative to humerus compared with other saurosphargids; 8) ilium with completely reduced iliac blade (unlike *Largocephalosaurus*, in which the iliac blade is reduced but present) (ilium morphology unknown in *Sinosaurosphargis*); 9) presence of a single distal tarsal (distal tarsal IV) (different from *Largocephalosaurus*, which possesses two distal tarsals – III and IV) (number of tarsals unknown in *Sinosaurosphargis*).

### Description and comparisons

HFUT YZSB-19-109 comprises three blocks (Figure 2). The first large block contains mostly disarticulated parts from the anterior right portion of the postcranial skeleton – two cervical neural arches and three cervical ribs, one dorsal centrum and two dorsal neural arches (all three preserved in articulation), nine dorsal ribs, one median, 11 lateral and 13 lateralmost gastral elements, a single osteoderm, the right scapula, right coracoid, right humerus and right radius (Figure 3). The second small block contains the distal ends of likely the last cervical rib and the six most anterior dorsal ribs, five lateralmost gastral elements, as many as seven osteoderms, a partial left coracoid and a left humerus, all preserved in ventral view (Figure 4). The third large block preserves the articulated posterior part of the body in ventral view, including six posterior dorsal vertebrae and associated ribs, three sacral vertebrae and ribs, an articulated series of 11 anterior caudals with associated ribs, chevrons and osteoderms, 14 lateralmost gastralia, a single large osteoderm, a partial right and a complete left pelvic girdle and a complete left hindlimb (Figures 5 and 6). Selected measurements of HFUT YZSB-19-109 are available in Supplementary Table 1.

**Figure 3.**
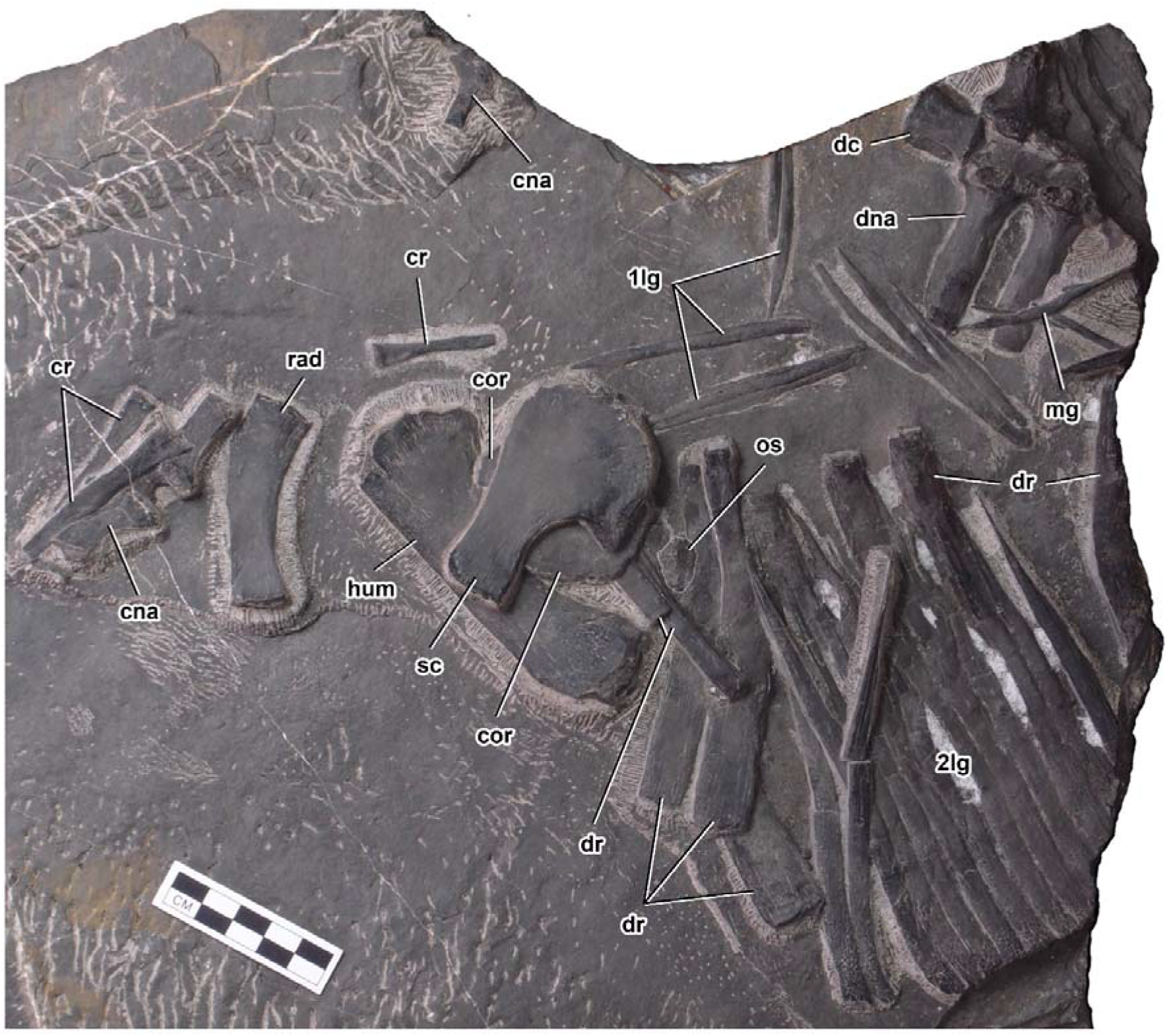
Anterior vertebral column, right pectoral girdle and right forelimb elements of *Prosaurosphargis yingzishanensis*. Abbreviations: 1lg, first lateral gastral element; 2lg, second lateral gastral element; can, cervical neural arch; cor, coracoid; cr, cervical rib; dc, dorsal centrum; dna, dorsal neural arch; dr, dorsal rib; hum, humerus; mg, median gastral element; os, osteoderm; rad, radius; sc, scapula. Scale bar = 5 cm.

**Figure 4.**
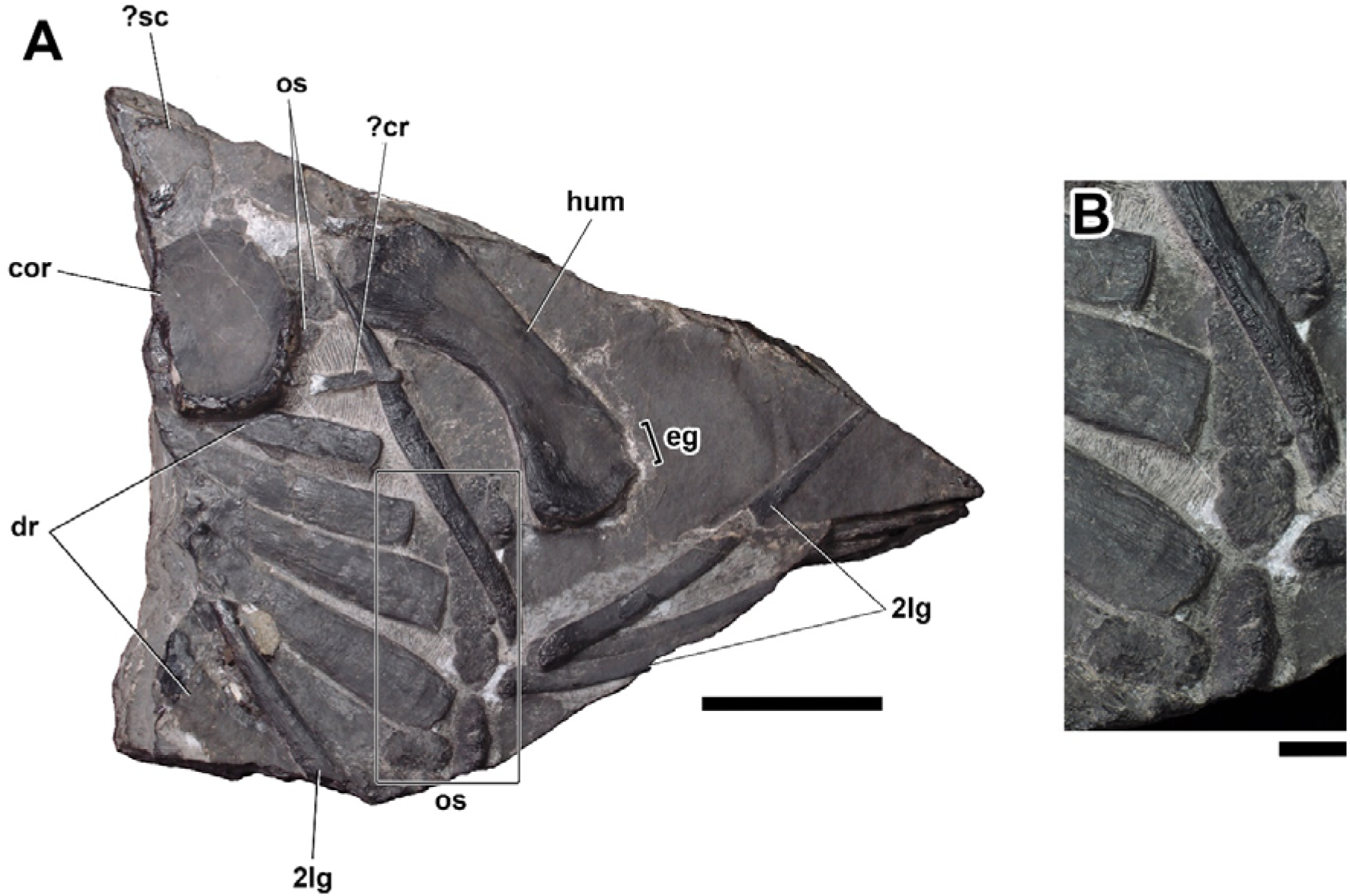
Dorsal ribs, lateral osteoderms, coracoid and humerus from the left side of the body of *Prosaurosphargis yingzishanensis* (A) and detail of lateral osteoderms (B). Abbreviations: 2lg, second lateral gastral element; cor, coracoid; ?cr, ?cervical rib; dr, dorsal rib; eg, ectepicondylar groove; hum, humerus; os, osteoderm, ?sc, ?scapula. Scale bar = 5 cm in (A) and 1 cm in (B).

**Figure 5.**
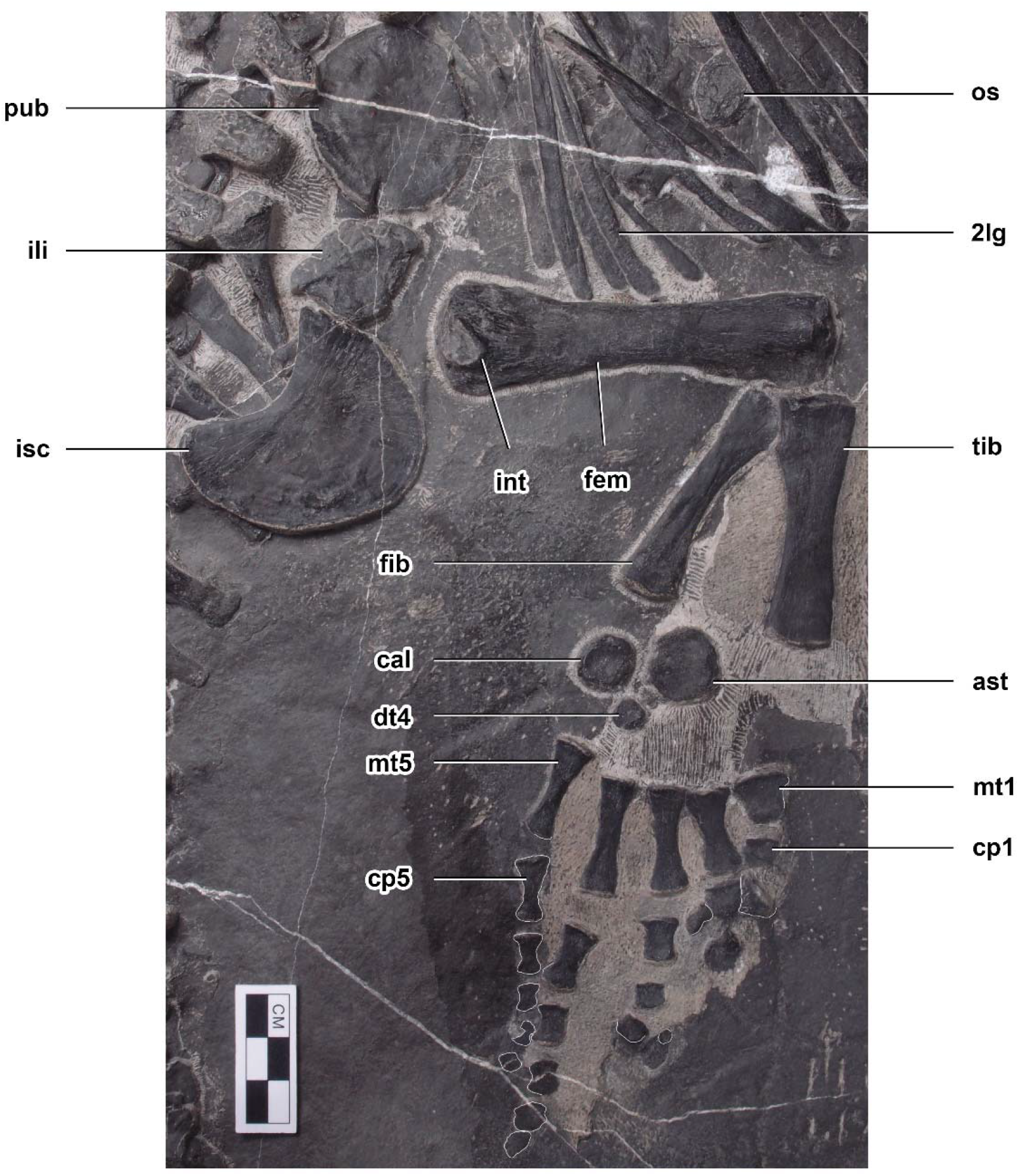
Pelvic girdle and hindlimb of *Prosaurosphargis yingzishanensis*. Abbreviations: 2lg, second lateral gastral element; ast, astragalus; cal, calcaneum; cp1, carpal I; cp5, carpal V; dt4, distal tarsal IV; fem, femur; fib, fibula; ili, ilium; int, internal trochanter; isc, ischium; mt1, metatarsal I; mt5, metatarsal V; os, osteoderm; pub, pubis; tib, tibia. Scale bar = 3 cm.

**Figure 6.**
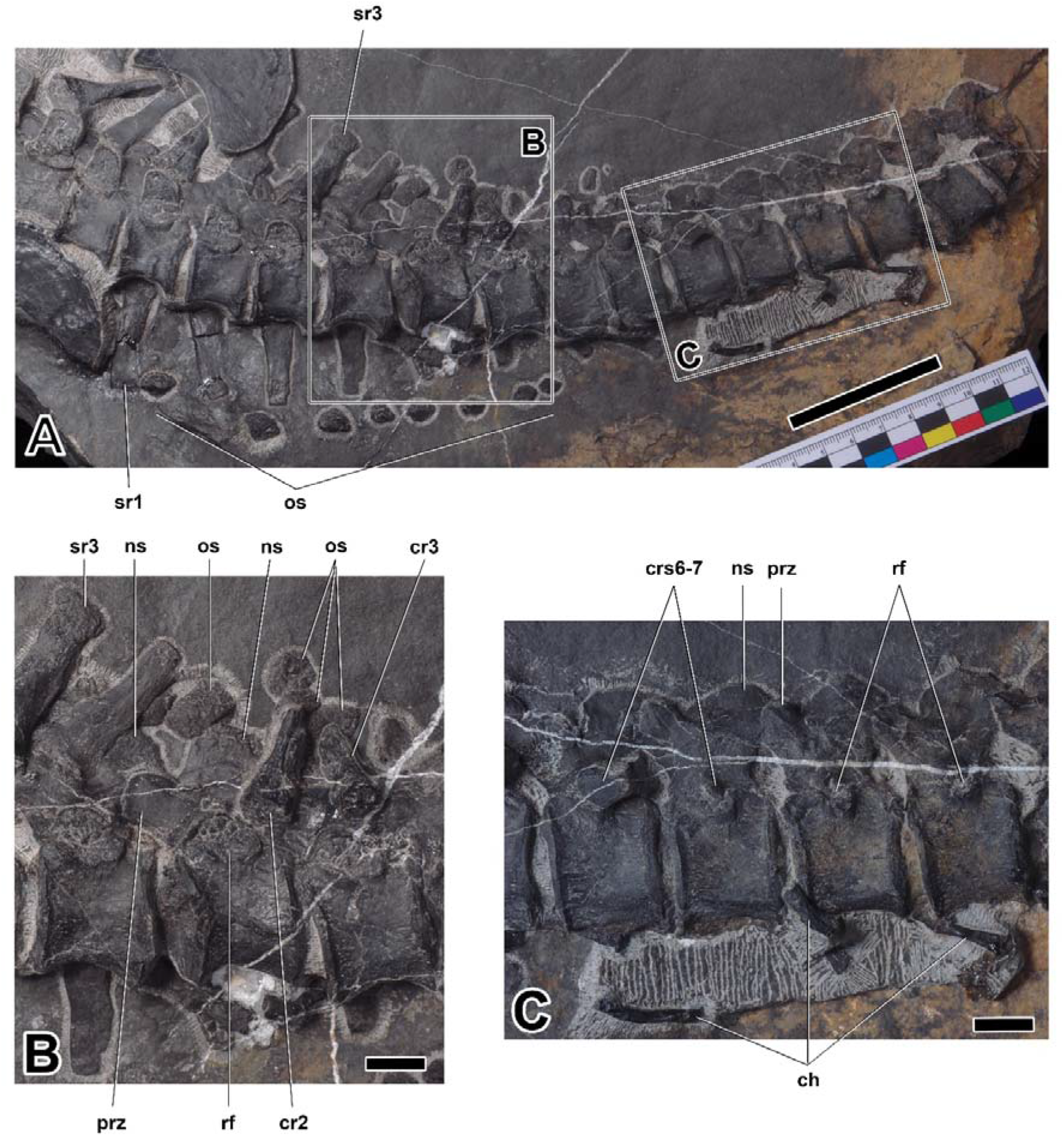
Sacral and caudal vertebrae of *Prosaurosphargis yingzishanensis*. Abbreviations: ch, chevron; cr2, second caudal rib; cr3, third caudal rib; crs6-7, sixth and seventh caudal ribs; ns, neural spine; os, osteoderm; prz, prezygapophysis; rf, rib facet; sr1, first sacral rib; sr3, third sacral rib. Scale bar = 5 cm in (A) and 1 cm in (B) and (C).

#### Axial skeleton

##### Vertebrae

The vertebral column of HFUT YZSB-19-109 is represented by two disarticulated cervical neural arches, a single anterior dorsal centrum and two anterior dorsal neural arches (all three preserved in articulation) and an articulated series of 20 vertebrae comprising six posterior dorsal neural arches, three complete sacral vertebrae and eleven complete caudal vertebrae.

Two cervical neural arches are preserved in HFUT YZSB-19-109 (Figure 3). Only the left side of a small neural arch, representing one of the anterior post-axial cervicals, is preserved in dorsal view, whereas a larger neural arch, belonging to one of the posterior cervicals, is completely preserved and exposed in ventral view, but it is partially obscured by an overlying cervical rib and is rotated by approximately 180 degrees, so that its anterior end faces posteriorly. The transverse processes of the cervical neural arches extend posterolaterally and are proportionally shorter than the transverse processes of the dorsal neural arches. A well-developed prezygapophysis is clearly visible in the smaller neural arch. Well-developed postzygapophyses are preserved in both the anterior and posterior cervical neural arches, where they are visible protruding from just underneath the overlying cervical rib in the latter. Approximately oval, anteroposteriorly elongate facets for articulation with the centrum are exposed in the posterior cervical neural arch.

The only preserved dorsal centrum is exposed in ventral view (Figure 3). It is mediolaterally constricted at its anteroposterior mid-length, resembling the dorsal centra of other saurosphargids (Huene 1936:taf. XIII-2; Li et al. 2011:fig. 3B; Li et al. 2014:fig. 5a). The pedicels of the neural arches are anteroposteriorly elongate and mediolaterally narrow, resembling those preserved in *Saurosphargis volzi* (Huene 1936:taf. XIII-2). The transverse processes are straight, mediolaterally elongated and anteroposteriorly expanded laterally, achieving their greatest width at approximately 2/3 of their mediolateral length. The maximum width of the transverse process is greater than the maximum width of the space between the transverse processes of adjacent neural arches (Figure 3), a condition similar to that exhibited by *Saurosphargis volzi* (Huene 1936:taf. XIII-2) and *Sinosaurosphargis yunguiensis* (Li et al. 2011:fig. 3B), but different from *Largocephalosaurus qianensis* (Li et al. 2014:fig. 2c) and *Eusaurosphargis dalsassoi* (Scheyer et al. 2017:fig. 6), in which the widths of the spaces between the dorsal transverse processes are approximately equal or greater than the widths of the transverse processes.

Five posterior dorsal neural arches and a single complete posterior dorsal vertebra are preserved in articulation in HFUT YZSB-19-109 and are exposed in ventral view (in adddition, a fragmentary centrum is associated with the second preserved posterior dorsal neural arch). The articulation surfaces for the centra located on the pedicels of the posterior dorsal neural arches are anteroposteriorly elongate but seem mediolaterally broader and not as well demarcated from the transverse processes as those in the more anterior dorsal neural arches. The posterior dorsal transverse processes gradually become mediolaterally shorter and anteroposteriorly broader posteriorly. Well-developed prezygapophysis-postzygapophysis articulations are preserved and exposed between the second and fifth posterior dorsal neural arches. Three complete sacral vertebrae are preserved and exposed in left ventrolateral view, as indicated by the associated sacral ribs with expanded distal ends (Figure 6A). A series of 11 articulated and complete caudal vertebrae is also preserved, with two anterior caudals exposed in left ventrolateral view and the remaining caudals exposed in lateral view (Figure 6). The sacral and caudal centra are amphicoelus, but in some cases their articular facets are obscured by matrix or adjacent centra, giving a false impression of procoely or opisthocoely. The caudal centra are dorsoventrally taller than anteroposteriorly long and gradually decrease in size posteriorly. The caudal neural arches bear well-developed, anterodorsally inclined prezygapophyses. The transverse processes of the caudal neural arches are mediolaterally short and anteroposteriorly broad. The neural spines of some caudal neural arches are exposed in lateral view (Figure 6B). They are dorsoventrally short, anteroposteriorly broad and possess a convex dorsal margin. A rugose dorsolateral surface, reminiscent of a similar surface present in the dorsal neural spines of *Helveticosaurus zollingeri* (Rieppel 1989a:fig. 6), *Augustasaurus hagdorni* (Sander et al. 1997:fig. 6) and *Pomolispondylus biani* (Cheng et al. 2022:fig. 4a) is preserved in some of the anterior caudal neural spines. Rib facets are visible from the second sacral to the penultimate preserved caudal centrum (Figure 6). The dorsal portions of the rib facets are located on the ventrolateral surfaces of the neural arches, whereas their ventral portions are located on the dorsolateral surfaces of the centra and are demarcated by a prominent, ventrally arcuate ridge (Figure 6B). The surface of the rib facets is rugose. The last visible minute caudal rib is preserved in articulation with the seventh caudal centrum, but small rib facets are also preserved in the following three centra (Figure 6C). It is not clear, however, if small ossified caudal ribs were associated with these centra in life.

##### Ribs

Four cervical ribs are preserved in HFUT YZSB-19-109 (Figures 3 and 4). They possess a single, expanded head and narrow distal ends. The preserved cervical ribs are approximately straight. HFUT YZSB-19-109 also preserves 18 dorsal ribs, but most of them are incomplete and/or obscured by overlying skeletal elements, such as gastralia or other ribs (Figures 3, 4 and 6A). The anterior dorsal ribs are much more robust than the cervical ribs, being proximodistally longer and anteroposteriorly broader (Figures 3 and 4). They have a single head and greatly expanded distal portions that abut against each other, forming a characteristic ‘rib-basket’ also present in other saurosphargids (Huene 1836; Li et al. 2011, 2014), but differ from *Eusaurosphargis dalsassoi*, in which the dorsal ribs are narrow and widely spaced (Scheyer et al. 2017). The dorsal ribs do not bear a distinct uncinate process, being similar in this respect to *Sinosaurosphargis yunguiensis* (Li et al. 2011:fig. 3b), but differ from *Largocephalosaurus, Saurosphargis volzi, Eusaurosphargis dalsassoi* and a possible isolated saurosphargid rib from the Late Triassic of Switzerland, all of which possess a more or less well-developed uncinate process (Li et al. 2014; Scheyer et al. 2017; Scheyer et al. 2022). Four posterior dorsal ribs are preserved in HFUT YZSB-19-109; they have an expanded head, a narrow distal end and are much shorter and slender in comparison with the more anterior dorsal ribs. As a consequence, they did not form part of the closed ‘rib-basket’. Three pairs of sacral ribs are preserved in HFUT YZSB-19-109, although the right sacral ribs are partially preserved as impressions (Figure 6A). They are proximodistally short and possess an expanded distal end. The second left sacral rib bears a small posteroproximal process. The caudal ribs are proximodistally short, possess a broad head and taper distally (Figure 6). The anterior caudal ribs in HFUT YZSB-19-109 are shorter than the sacral ribs, in contrast to *Largocephalosaurus qianensis*, in which the anterior caudal ribs are longer than the sacral ribs (Li et al. 2014:fig. 5d). The caudal ribs decrease in size posteriorly and extended at least to the level of the seventh caudal centrum, although it is not clear if caudal ribs extended posteriorly beyond the seventh caudal (see above) (in *Largocephalosaurus qianensis* the caudal ribs extend to the level of the 10th or 11th caudal centrum; Li et al. 2014).

##### Chevrons

Eight chevrons are preserved in HFUT YZSB-19-109 in association with some of the posterior caudal vertebrae (Figure 6A, C). In lateral view, the chevrons are slender and straight or display a gentle ventral curvature. The chevrons have a proximal end that is approximately equal in size or only slightly expanded relative to the distal end.

##### Gastralia

The gastral rib basket is partially preserved in HFUT YZSB-19-109 (Figures 3–5). Like in *Placodus* and eosauropterygians (Drevermann 1933; Rieppel 1989b; Shang et al. 2011), each gastral rib was composed of five elements – a median element, two lateral (first lateral) elements and two lateralmost (second lateral) elements. In *Sinosaurosphargis*, the gastral ribs were described as comprising three elements – one median and two lateral elements, whereas the number of gastral rib elements was not specified for *Largocephalosaurus* (Li et al. 2011, 2014). A single anterior median gastral element, eleven lateral and thirty-three lateralmost elements are preserved in HFUT YZSB-19-109. The median element is weakly angulated posteriorly, whereas the first lateral elements form proportionally short rods with a blunt medial end and a pointed lateral end (Figure 3) and closely resemble the first lateral gastral elements of *Placodus* (Drevermann 1933). The lateralmost elements are concentrated into two articulated series comprising thirteen (anterior block; Figure 3) and fourteen (posterior block; Figure 5) elements each. They form mediolaterally elongate rods which are approximately straight or gently angulated posteriorly. The medial end of the lateral element is narrow and tapers into a pointed apex, whereas the lateral end is broadened and blunt with a distinctly rugose/granulated surface. One anteriorly positioned first lateral element bifurcates laterally, forming two lateral prongs (Figure 3).

#### Osteoderms

Several osteoderms are preserved in HFUT YZSB-19-109. A single osteoderm is preserved in the anterior block between the anterior dorsal ribs (Figure 3) and a second isolated osteoderm is preserved in the posterior block between the posterior gastral elements (Figure 5). These osteoderms are approximately oval in outline and possibly represent osteoderms forming parasaggital rows similar to those in *Largocephalosaurus* (Li et al. 2014; Scheyer et al. 2022), but they could also represent displaced lateral osteoderms. A partial series of as many as seven osteoderms is preserved between the left coracoid and humerus and along the distal ends of a series of partially preserved left dorsal ribs (Figure 4). These osteoderms represent the left lateral osteoderm row and are anteroposteriorly elongate, have an irregular, sub-oval or sub-rectangular outline and a densely pitted surface. The most posterior of these osteoderms bears a prominent ridge extending along the midline of its exposed surface. In all these features, these osteoderms resemble the lateral osteoderms reported for *Largocephalosaurus polycarpon* (Li et al. 2014:fig. 4h).

Another partial series of small osteoderms is preserved lateral to the distal ends of the right sacral and anterior caudal ribs, indicating that the lateral osteoderm rows extended to the level of the seventh caudal vertebra (Figure 6A). A few small osteoderms are also preserved in close association with the anterior caudal neural spines and seem to have formed a median row along the dorsal midline (Figure 6B), likely overlying the caudal neural spines in a manner similar to that in *Largocephalosaurus* (Li et al. 2014) and *Placodus inexpectatus* (Jiang et al. 2008). In *Largocephalosaurus qianensis*, a dense covering of small osteoderms is present on the dorsal surface of the neck, trunk and caudal region (Li et al. 2014), but no such osteoderms were found in association with HFUT YSZB-19-109. A dense covering of osteoderms is also absent in *Largocephalosaurus polycarpon* (Cheng et al. 2012; Li et al. 2014).

#### Pectoral girdle

##### Scapula

The right scapula is completely preserved in HFUT YZSB-19-109 (Figure 3). In addition, a large and broad broken bone fragment likely representing the left scapula is visible lying anterodorsally to the left coracoid (Figure 4). The scapula possesses an anteroproximally expanded glenoid portion and a much narrower, straight and posterodorsally projecting scapular blade. The glenoid portion forms a low acromion process proximodorsally. The coracoid facet is relatively broad and convex, whereas the glenoid facet is straight and short and oriented nearly parallel to the long axis of the scapular blade. The scapular blade is separated from the glenoid portion by a deep posteroventral notch. The scapular blade is straight, with gently concave anterodorsal and posteroventral margins and an approximately straight posterior margin. In general shape and proportions, the scapula of HFUT YZSB-19-109 closely resembles that of *Largocephalosaurus* (Cheng et al. 2012:fig. 3; Li et al. 2014:fig 4d), *Eusaurosphargis dalsassoi* (Nosotti and Rieppel 2003:fig. 18; Scheyer et al. 2017:fig. 3), and *Corosaurus alcovensis* (Storrs 1991:fig. 12), although the notch separating the glenoid portion from the scapular blade is much deeper and narrower in the latter.

##### Coracoid

The coracoids are only partially visible in HFUT YZSB-19-109 – the right coracoid is almost entirely covered by the right scapula (Figure 3), whereas the left coracoid is broken proximomedially (Figure 4). The preserved parts of both coracoids indicate that it was a dorsoventrally flat, plate-like element, approximately sub-oval in outline with a rounded distal margin, like in *Largocephalosaurs qianensis* (Li et al. 2014:fig. 5a), *Placodus gigas* (Drevermann 1933:pls 12, 14), *Majiashanosaurus discocoracoidis* (Jiang et al. 2014:fig. 3A), ‘*Lariosaurus*’ *sanxiaensis* (Li and Liu 2020:fig. 3d–f), a referred specimen of *Hanosaurus hupehensis* (Wang et al. 2022:fig. 1G–H) and possibly *Saurosphargis volzi* (a partially exposed, putative coracoid is preserved on the ventral surface of the only figured part of the type specimen; see Huene 1936:pl. XIII and Nosotti and Rieppel 2003:fig 11). It slightly differs, however, from the coracoid of *Helveticosaurus zollingeri* (Rieppel 1989:fig. 6) and *Eusaurosphargis dalsassoi* (Nosotti and Rieppel 2003:fig. 18; Scheyer et al. 2017:fig. 6), in which the anterior margin of the coracoid is weakly concave and the anterodistal corner of the bone forms an approximately right angle. A small notch in the proximal part of the right coracoid, seen just above the anterior margin of the right scapula, likely represents the coracoid foramen. The exposed ventral surface of the left coracoid bears numerous radial striations extending from the centre of the bone towards its outer margins.

#### Forelimb

##### Humerus

Both humeri are preserved in HFUT YZSB-19-109 – the right humerus is complete, but partially overlapped by the right scapular blade (Figure 3), whereas the left humerus is slightly broken proximally (Figure 4). The humerus is proximodistally elongate and posteriorly curved along its proximodistal axis, like in other saurosphargids (Li et al. 2011, 2014; Cheng et al. 2012), placodonts (Rieppel 2000a), *Majiashanosaurus discocoracoidis* (Jiang et al. 2014), ‘*Lariosaurus*’ *sanxiaensis* (Li and Liu 2020), a referred specimen of *Hanosaurus hupehensis* (Wang et al. 2022) and numerous eosauropterygians (Rieppel 1994). However, in contrast to *Largocephalosaurus*, the anterior margin of the humerus is not convex, but straight, making the humerus more similar to that of *Placodus* and *Nothosaurus* (Rieppel 1994) (shape of anterior margin of humerus unknown in *Sinosaurosphargis*). The posterior margin of the humerus is concave. The proximal and distal ends of the humerus are expanded, but a distinct humeral head and distal condyles are not present. Anteriorly, the distal end of the humerus bears a shallow ectepicondylar groove (notch), like in *Largocephalosaurus qianensis* (Li et al. 2014:fig. 6b), but unlike *Largocephalosaurus polycarpon* (Cheng et al. 2012:fig. 3), which possesses an entepicondylar foramen. Anteroproximally, the exposed surface of the right humerus bears a short, straight groove, which likely represents a muscle/tendon origin/insertion site.

##### Radius

A disarticulated right radius is completely preserved in HFUT YZSB-19-109 in close proximity to the right humerus and right pectoral girdle (Figure 3). The radius is proximodistally elongate and anteriorly curved, with a convex anterior margin and a concave posterior margin. The proximal and distal ends of the radius are straight and slightly expanded, with the proximal end being anteroposteriorly broader than the distal end. The radius is similar in general shape to that in other saurosphargids (Li et al. 2011:fig. 1; Cheng et al. 2012:fig. 3; Li et al. 2014:fig. 6b), *Majiashanosaurus discocoracoidis* (Jiang et al. 2014:fig. 4), *Placodus inexpectatus* (Jiang et al. 2008:fig. 3A) and *Helveticosaurus zollingeri* (Rieppel 1989:fig. 6), but differs from the straight and anteriorly and posteriorly concave radius of *Eusaurosphargis dalsassoi* (Scheyer et al. 2017:fig. 6). The radius/humerus proximodistal length ratio in HFUT YZSB-19-109 is ∼0.55, in line with *Eusaurosphargis dalsassoi* (∼0.51–0.57; Scheyer et al. 2017) and *Placodus inexpectatus* (∼0.56; Jiang et al. 2008), but is significantly smaller than the ratio in *Sinosaurosphargis yunguiensis* (∼0.66; Li et al. 2011), *Largocephalosaurus polycarpon* (∼0.74; Cheng et al. 2012), *Largocephalosaurus qianensis* (∼0.67–0.72; Li et al. 2014), a referred specimen of *Hanosaurus hupehensis* (∼0.67; Wang et al. 2022) and *Majiashanosaurus discocoracoidis* (∼0.74; Jiang et al. 2014). This indicates that HFUT YZSB-19-109 had a proportionally shorter forearm compared with other saurosphargids and eosauropterygians.

#### Pelvic girdle

##### Ilium

The left ilium is completely preserved in HFUT YZSB-19-109 and is exposed in lateral view (Figure 5). The ilium is approximately pentagonal in outline and anteroposteriorly expanded distally. It consists entirely of an acetabular portion and bears no distinct iliac blade. This is in stark contrast to *Largocephalosaurus qianensis* (Li et al. 2014:fig. 6a, d), *Eusaurosphargis dalsassoi* (Scheyer et al. 2017:fig. 3b), *Placodus gigas* (Rieppel 1995:fig. 41) and *Corosaurus alcovensis* (Storrs 1991:fig. 15), which possess ilia with a well-developed, posteriorly projecting iliac blade. The ilium of HFUT YZSB-19-109 is more similar to the ilia of *Sanchiaosaurus dengi* (Rieppel 1999:fig. 8H), *Diandongosaurus acutidentatus* (Liu et al. 2015:fig. 1), *Neusticosaurus edwardsii* (Carroll and Gaskill 1985:fig. 5i) and *Lariosaurus buzzi* (Tschanz 1989:text-fig. 6h, i), in which the iliac blade is strongly reduced, but is still present as a small, posterodorsally projecting process.

##### Pubis

The left pubis is completely preserved in HFUT YZSB-19-109 and exposed in ventral view, whereas only the ventral surface of the distal portion of the right pubis is preserved (Figure 5). The pubis is approximately oval in outline, being proximodistally longer than anteroposteriorly broad and bears a proximally positioned, open obturator foramen. The pubis of HFUT YZSB-19-109 resembles the pubis of *Largocephalosaurus qianensis* (Li et al. 2014:fig. 5d) in outline and proportions. It is also similar to the pubis of the holotype and referred specimens of *Hanosaurus hupehensis* (Rieppel 1998a:fig. 5; Wang et al. 2022:fig. 1I) and *Pararcus diepenbroekii* (Klein and Scheyer 2013:fig. 5), which are however more circular in outline, being approximately as wide proximodistally as long anteroposteriorly. The pubis of HFUT YZSB-19-109 differs markedly from the anteriorly and posteriorly shallowly emarginated pubis of *Eusaurosphargis dalsassoi* (Scheyer et al. 2017:fig. 8) and *Placodus gigas* (Drevermann 1933:pl. 13) and the deeply emarginated pubis of eosauropterygians (e.g. Rieppel 2000a).

##### Ischium

The left ischium of HFUT YZSB-19-109 is completely preserved, but is rotated 180° relative to its life position and is exposed in dorsal view, whereas only the distal portion of the right ischium is preserved (Figure 5). The ischium forms an anterodistally convex and posteroproximally concave plate, similar to that of *Largocephalosaurus qianensis* (Li et al. 2014:fig. 5d), *Pararcus diepenbroekii* (Klein and Scheyer 2013) and *Hanosaurus hupehensis* (Rieppel 1998a; Wang et al. 2022: fig. 1I), but differs markedly from the anteriorly emarginated ischium of *Eusaurosphargis dalsassoi* (Scheyer et al. 2017), the posterodistally emarginated ischium of *Henodus chelyops* (Huene 1836), and the anteriorly and posteriorly emarginated ischia of some other placodonts and eosauropterygians (Rieppel 2000a).

#### Hindlimb

Femur: The left hindlimb is completely preserved in HFUT YZSB-19-109. The femur is exposed in posterior view (Figure 5). The shaft of the femur is straight and both the proximal and distal ends are expanded. The internal trochanter is well-developed and located proximally, as in *Largocephalosaurus qianensis* (Li et al. 2014:fig. 6d), *Simosaurus gallardoti* and *Nothosaurus* (Rieppel 1994:figs 33, 62), but differs from the more distally located trochanter in the holotype of *Hanosaurus hupehensis* (Rieppel 1998a:fig. 4c). Distally, the femur produces weakly-developed but still distinct condyles for the tibia and fibula, separated ventrally by a shallow popliteal area. The humerus/femur ratio in HFUT YZSB-19-109 is ∼1.19, which is similar to the ratio in *Largocephalosaurus qianensis* (∼1.20; Li et al. 2014), but is significantly greater than the ratio in *Helveticosaurus zollingeri* (∼1.11; Rieppel 1989), *Placodus inexpectatus* (∼1.05; Jiang et al. 2008) and *Eusaurosphargis dalsassoi* (∼0.89; Scheyer et al. 2017).

##### Tibia and fibula

The tibia is proximodistally slightly longer than the fibula and possesses proximally and distally expanded ends, with the proximal end slightly broader than the distal end (Figure 5). Posteroproximally, the tibia bears a proximodistally elongate, shallow facet for the fibula. The anterior and posterior margins of the tibia are gently concave. The fibula possesses an approximately straight posterior margin and a concave anterior margin (Figure 5). The proximal and distal ends of the fibula are slightly expanded, with the distal end being slightly broader than the proximal end. This is in contrast to *Largocephalosaurus qianensis*, in which the proximal end of the fibula is markedly broader than the distal end (Li et al. 2014:fig. 6d).

##### Tarsals

Three tarsals are present in the left hindlimb of HFUT YZSB-19-109 – the largest one being the astragalus, the medium-sized representing the calcaneum, and the smallest element interpreted here as distal tarsal IV (Figure 5). The tarsals are sub-circular in outline, with the astragalus possessing minute notches anteriorly and posteriorly, and their ventral surfaces are weakly concave. The morphology of the tarsals of HFUT YZSB-19-109 is similar to that of *Largocephalosaurus*, which, however, possesses 4 tarsals instead of 3 (astragalus, calcaneum and distal tarsals III and IV; Li et al. 2014:fig. 5d).

##### Metatarsals

All five metatarsals are preserved in the left hindlimb of HFUT YZSB-19-109 (Figure 5). Metatarsal I is the proximodistally shortest metatarsal, being much broader proximally than distally. It is also the most robust of the metatarsals. Metatarsals II–V are slender, approximately cylindrical in outline, with expaned proximal and distal ends. The proximodistal length of the metatarsals increases from metatarsal II–IV, with metatarsal IV being the longest of all metatarsals (similar to *Largocephalosaurus polycarpon* but different from *Largocephalosaurus qianensis*, in which metatarsal III is the longest; Li et al. 2014); the length of metatarsal V is comparable to that of metatarsal III. Metatarsal V is proportionally the most slender of the metatarsals (similar to the condition in *Largocephalosaurus qianensis*, but unlike *Largocephalosaurus polycarpon*, in which metatarsal IV is the most slender; Li et al. 2014).

##### Phalanges

The pedal phalanges are completely preserved in the left pes of HFUT YZSB-19-109 (Figure 5). The proximal phalanges of digits 1 and 2 are rectangular in outline, whereas those of digits 3–5 are cylindrical and have expanded proximal and distal ends. Distally, the carpals become proximodistally shorter and sub-rectangular in outline. The phalangeal formula is 2-3-4-5-5 (unguals 2 and 3 are slightly displaced).

### Ontogeny and body size

Articulated anterior and posterior dorsal neural arches preserved without their respective centra indicate a weak connection between these elements in HFUT YZSB-19-109. This demonstrates that the specimen likely represents an osteologically immature individual that has not reached its full adult body size. Based on the humerus:total body length and femur:total body length ratios of the type specimen of *Largocephalosaurus qianensis* (specimen IVPP V 15638, total body length = 2317 mm; Li et al. 2014), the only completely preserved saurosphargid specimen discovered to date, the total length of HFUT YZSB-19-109 is estimated to have reached between 1468 mm to 1583 mm, respectively. However, due to the limited availability of comparative material, differences in body proportions between the known saurosphargids (see above) and the incomplete nature of HFUT YZSB-19-109, the provided total body length estimate should be treated with caution.

### Phylogenetic results

The phylogenetic analysis recovered 48 most parsimonious trees (MPTs) of 1010 steps each (CI = 0.271, RI = 0.625) (Figure 7). *Prosaurosphargis* was recovered as a member of Saurosphargidae, forming a ploytomy with *Sinosaurosphargis* and *Largocephalosaurus*. Saurosphargidae is supported by the following six unambiguous synapomorphies: caudal lateral projections (transverse processes) beyond fifth caudal present (char. 126.1); distinct free anterior process of cervical ribs absent (char. 129.0); dorsal ribs transversely broadened and in antero-posterior contact with each other, forming closed ‘rib-basket’ (char. 135.1); distal width of haemal spine equivalent to proximal width (char. 137.0); lateral gastralia expanded and flat (char. 140.1); distal end of ulna distinctly expanded (char. 167.1).

**Figure 7.**
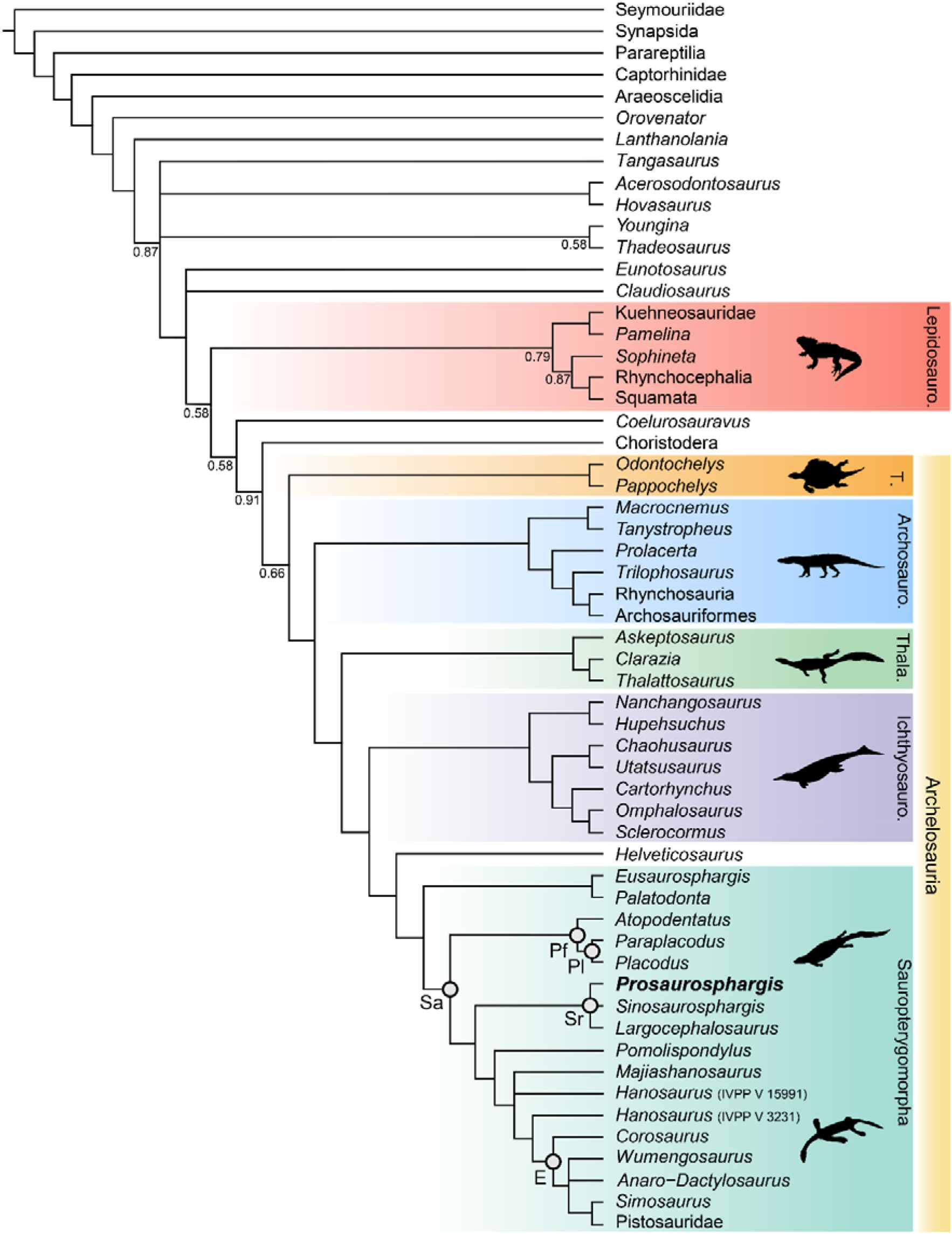
Phylogenetic relationships of *Prosaurosphargis yingzishanensis* within Diapsida. The 50% majority rule consensus of 48 most parsimonious trees (MPTs) obtained from analysis of the updated dataset of Qiao et al. (2022). Numbers below nodes indicate proportion of MPTs that recover that node if it is lower than 1. Abbreviations: Archosauro., Archosauromorpha; E, Eosauropterygia; Ichthyosauro., Ichthyosauromorpha; Lepidosauro., Lepidosauromorpha; Pf, Placodontiformes; Pl, Placodontia; Sa, Sauropterygia; Sr, Saurosphargidae; T., Testudinata; Thala., Thalattosauria. Silhouettes from phylopic.org.

Saurosphargidae was recovered within Sauropterygia, as the sister-group to the linage leading to Eosauropterygia. The saurosphargid-eosauropterygian clade is supported by six unambiguous synapomorphies: frontal, butterfly-shaped with antero- and postero-lateral processes absent (ch. 29.0); long postorbital posterior process contacting squamosal (ch. 32.0); mandibular articulations displaced to a level distinctly behind occipital condyle (ch. 87.1); neural canal evenly proportioned (ch. 122.0); three sacral ribs (ch. 130.1); total number of carpal ossifications more than three (ch. 194.0).

*Pomolispondylus*, recently proposed as a sister-taxon of Saurosphargidae (Cheng et al. 2022), was recovered as the most basal member of the grade leading to Eosauropterygia (*Pomolispondylus* + eosauropterygian lineage supported by a single unambiguous synapomorphy: transverse processes of neural arches of the dorsal region relatively short [ch. 124.0]). The type (Rieppel 1998a) and referred (Wang et al. 2022) specimens of *Hanosaurus* were not recovered in a monophyletic group – both specimens were recovered alongside *Majiashanosaurus* in a polytomy at the base of Eosauropterygia (clade comprising *Hanosaurus, Majiashanosaurus* and Eosauropterygia supported by a single unambiguous synapomorhpy: osteoderms absent [ch. 143.0]). *Corosaurus* was recovered as the earliest-diverging member of Eosauropterygia, whereas *Wumengosaurus* was recovered outside of the clade comprising pachypleurosaurs, nothosaurs and pistosaurs.

The herbivorous sauropterygian *Atopodentatus* was recovered as the sister-taxon of placodonts (represented in our dataset by *Paraplacodus* and *Placodus*) within Placodontiformes, supported by four unambiguous synapomorphies: contact of the prefrontal and postfrontal excludes frontal from dorsal orbital margin (char. 21.1); interpterygoid vacuity absent (char. 81.1); spelnial enters mandibular symphysis (char. 88.0); and femur internal trochanter well developed (char. 203.0).

*Palatodonta* was recovered as the sister-taxon of *Eusaurosphargis*. This clade is supported by one unambiguous synapomorphy – a small premaxilla (char. 196.1). The clade comprising *Palatodonta* + *Eusaurosphargis* was recovered as the sister-group to Sauropterygia within Sauropterygomorpha tax. nov. (see above). *Helveticosaurus* was recovered as the sister-group of Sauropterygomorpha; the clade comprising *Helveticosaurus* + Sauropterygomorpha is supported by the following six unambiguous synapomorphies: preorbital and postorbital regions of skull of subequal length (ch. 1.0); transverse processes of neural arches of the dorsal region distinctly elongated (ch. 124.1); scapula with a constriction separating a ventral glenoidal portion from a posteriorly directed dorsal wing (ch. 154.2); distal tarsal I absent (ch. 184.1); total number of tarsal ossifications less than four (ch. 192.1); total number of carpal ossifications two (char. 194.2).

Within Diapsida, the clade comprising *Helveticosaurus* + Sauropterygomorpha was recovered forming a clade with Ichthyosauromorpha, Thalattosauria and Archosauromorpha, a result similar to that recovered in some other recent broad-scale analyses of diapsid phylogenetic interrelationships (Neenan et al. 2015; Chen et al. 2014; Martínez et al. 2021). The three major marine reptile clades, Archosauromorpha and Testudinata were recovered within a monophyletic Archelosauria supported by four unambiguous synapomorphies – frontal with distinct posterolateral processes (ch. 26.1), frontal anterior margins oblique, forming an angle of at least 30 degrees with long axis of the skull (ch. 27.1), interclavicle anterior process or triangle conspicuously present (ch. 157.0) and upper temporal fossae present and distinctly smaller than the orbit (ch. 207.3).

## Discussion

### Marine reptile diversity of the Early Triassic Nanzhang-Yuan’an Fauna

*Prosaurosphargis* represents the stratigraphically oldest occurrence of Saurosphargidae, extending their fossil record back by approximately 4.5 Ma from the Middle (Pelsonian) to the Early (Olenekian) Triassic (Figure 8). Saurosphargids are thus the fourth major marine reptile lineage known from the Early Triassic Nanzhang-Yuan’an fauna, which also includes as many as seven species of hupehsuchians (Chen et al. 2015; Qiao et al. 2020), one species of ichthyosauriform (Chen et al. 2013), and as many as five taxa representing the sauropterygian lineage leading to Eosauropterygia (Young 1965; Rieppel 1998a; Li and Liu 2020; Cheng et al. 2022; Wang et al. 2022). Measuring approximately 1.5 m in total body length, *Prosaurosphargis* is one of the larger marine reptiles known from this ecosystem (despite the fact that the holotype likely represents an osteologically immature individual; see above), smaller only than an unnamed eosauropterygian (but possibly representing the same taxon as ‘*Lariosaurus*’ *sanxiaensis*/*Hanosaurus hupehensis* [referred specimen]) (body length of 3–4 meters; Chen et al. 2014b, 2016) and an indeterminate hupehsuchian (body length of ∼2.3 meters; Qiao et al. 2020). This indicates that the ecological capacity of the Nanzhang-Yuan’an shallow marine Early Triassic ecosystem to sustain a broad diversity of marine reptiles was even greater than previously thought (Li and Liu 2020).

**Figure 8.**
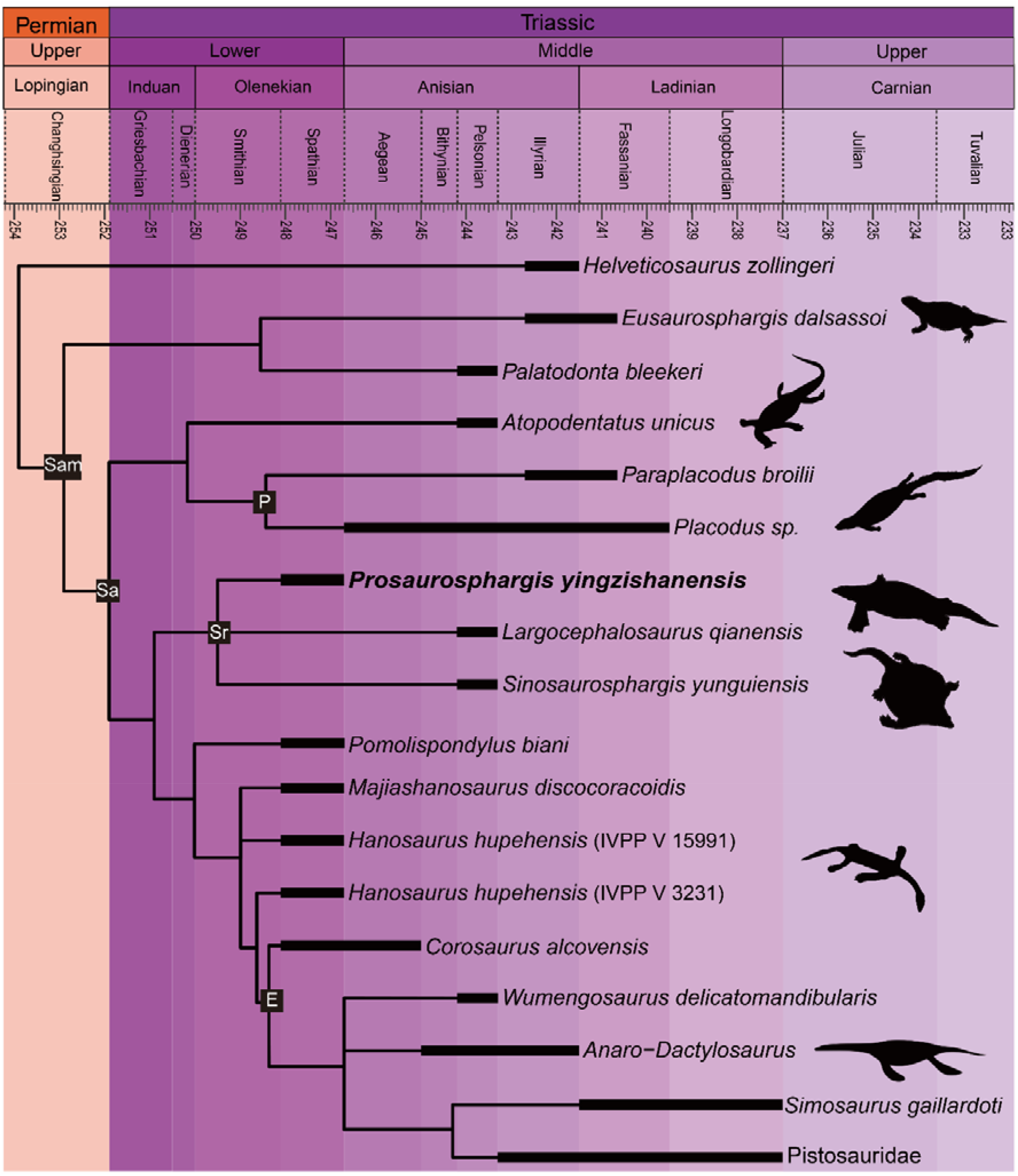
Time-scaled phylogenetic tree of Sauropterygomorpha. Abbreviations: E, Eosauropterygia; Pf, Placodontiformes; Pl, Placodontia; Sa, Sauropterygia; Sam, Sauropterygomorpha; Sr, Saurosphargidae. Silhouettes for *Atopodentatus, Paraplacodus* and eosauropterygians from phylopic.org.

### Phylogeny of Sauropterygomorpha

A clade comprising *Palatodonta* + *Eusarosphargis* and the lineage leading to *Helveticosaurus* are recovered as successive outgroups to Sauropterygia, with *Palatodonta* + *Eusaurosphargis* and Sauropterygia united within Sauropterygomorpha tax. nov. (see above). The status of *Eusaurosphargis* + *Palatodonta* as the sister-group to Sauropterygia is corroborated by the presence in *Eusaurosphargis* of morphological features otherwise known exclusively in sauropterygians: a clavicle applied to the medial surface of the scapula and a pectoral fenestration (Scheyer et al. 2017). Furthermore, *Eusaurosphargis* and Sauropterygia also share a similar foot morphology with metatarsal I being proximodistally much shorter than metatarsal IV and metatarsal V being long and slender (see above). *Helveticosaurus* shares the presence of a skull with preorbital and postorbital regions subequal in length with *Eusaurosphargis* and *Palatodonta*, the presence of elongated dorsal neural spines with *Eusaurosphargis*, placodonts and saurosphargids and the presence of a scapular constriction with *Eusaurosphargis* and sauropterygians. However, other anatomical features uniting it with Sauropterygomorpha include details of carpal and tarsal anatomy, which likely represent aquatic adaptations that might have evolved convergently in *Helveticosaurus* and aquatic representatives of Sauropterygomorpha (Chen et al. 2014a). Furthermore, *Helveticosaurus* lacks osteoderms, a feature present in early members of all major lineages within Sauopterygomorpha. Consequently, we interpret *Helveticosaurus* as a representative of a lineage closely related to Sauropterygomorpha but lying outside of it, that likely convergently evolved an aquatic lifestyle.

Our phylogenetic analysis does not recover *Palatodonta* as a sister-taxon of placodonts within Placodontiformes as was previously proposed (Neenan et al. 2013). Instead, *Palatodonta* is recovered as the sister-taxon of *Eusaurosphargis*. The *Palatodonta* + *Eusaurosphargis* clade recovered by our phylogenetic analysis is supported by one unambiguous synapomorphy – a small premaxilla (ch. 196.0). In addition, both taxa also share anteriorly positioned external nares (ch. 197.0). Both of these character states are considered as typical for terrestrial taxa and contrast with the enlarged premaxillae and posteriory displaced external narial openings characteristic for marine reptiles (Chen et al. 2014a). A terrestrial lifetyle was previously proposed for *Eusaurosphargis* on the basis of manus and pes anatomy and bone histology (Scheyer et al. 2017) and the femur of *Eusaurosphargis* is longer than its humerus (humerus:femur ratio ∼0.89; Scheyer et al. 2017), which indicates hindlimb dominance characteristic of a terrestrial lifestyle (Motani and Vermeij 2021). All this evidence strongly suggests that *Palatodonta* and *Eusaurosphargis* were terrestrial reptiles, likely representing the morphology of the last common terrestrial ancestor of Sauropterygia. *Palatodonta* is known from a single isolated skull (Neenan et al. 2013), whereas *Eusaurosphargis* is represented by two specimens with well preserved and largely complete postcrania, but only partially preserved skulls (Nosotti and Rieppel 2003; Scheyer et al. 2017). Because postcranial remains referable to *Eusaurosphargis* were also reported from the type locality of *Palatodonta*, including a *Palatodonta*-like dentary preserved in close association with typical *Eusaurosphargis*-like vertebrae (Scheyer et al. 2019; Willemse et al. 2019), it is very likely that *Palatodonta* is a junior synonym of *Eusaurosphargis*, but the discovery of well-preserved skulls with associated postcranial elements of both of these taxa are needed to further test this hypothesis.

*Atopodentatus* and placodonts are recovered within Placodontiformes and this grouping is supported by four unambiguous synapomorphies (see above). *Atopodentatus* and the early-diverging placodonts *Placodus* and *Paraplacodus* possess a humerus which is longer than the femur, indicating a high level of adaptation to an aquatic lifestyle (Motani and Vermeij 2021), but they also possess a massive femoral fourth trochanter and an ilium with a well-developed iliac blade (Jiang et al. 2008; Cheng et al. 2014a). These features indicate that the hindlimbs in both *Atopodentatus* and placodonts were likely still important in locomotion at the bottom of the sea floor and/or on shore in a marginal marine environment and suggest a slightly lower degree of adaptation to an aquatic lifestyle in placodontiforms than in saurosphargids and eosauropterygians, in which the fourth trochanter and iliac blade are more reduced (Rieppel 2000a; Li et al. 2014). The sister-group relationships of *Atopodentatus* and placodonts might also explain the absence of placodonts in Early Triassic fossil horizons worldwide. It is possible that the lineage leading to placodonts was represented in the Early Triassic by reptiles morphologically more similar to *Atopodentatus* than to placodonts and that the specialised placodont body plan did not evolve until the Middle Triassic. New discoveries of Early Triassic sauropterygians are likely to introduce new morphological data needed to test this hypothesis.

*Pomolispondylus* is not recovered as a sister-taxon of Saurosphargidae within Saurosphargiformes (contra Cheng et al. 2022), but as the most basal member of a grade of sauropterygians leading to Eosauropterygia. Such a phylogenetic position is supported by the presence in *Pomolispoindylus* of dorsal transverse processes that are relatively short mediolaterally and broad anteroposteriorly, more similar in proportions to the dorsal transverse processes of ‘*Lariosaurus*’ *sanxiaensis* (Li and Liu 2020:fig. 2) and *Hanosaurus hupehensis* (Wang et al. 2022:fig. 4g) – two other representatives of the grade leading to Eosauropterygia – than to the mediolaterally broad and anteroposteriorly narrow dorsal neural spines of saurosphargids (Figure 3; Li et al. 2011:fig. 3B; Li et al. 2014:fig. 6A). Furthermore, *Pomolispondylus* possesses rows of rudimentary osteoderms on its body flanks, which are much more reduced than those present in *Eusaurosphargis*, early-diverging placodonts and saurosphargids (Klein and Scheyer 2013; Li et al. 2014; Scheyer et al. 2017). Therefore, the osteoderms in *Pomolispondylus* likely represent a late stage of osteoderm reduction in the lineage leading to Eosauropterygia, rather than the first stages of osteoderm development in the saurosphargid lineage.

The type (Rieppel 1998a) and referred (Wang et al. 2022) specimens of *Hanosaurus* (Wang et al. 2022) and *Majiashanosaurus* (Jiang et al. 2014) are also recovered in the paraphyletic grade leading to Eosauropterygia, but in a position more derived than *Pomolispondylus*. This result is in contrast to previous studies which recovered these taxa as either the outgroup to Saurosphargidae + Sauropterygia (*Hanosaurus*; Wang et al. 2022), pachypleurosaurs (Rieppel 1998a; Neenan et al. 2015; Lin et al. 2021) or successive outgroups to a clade comprising Pachypleurosauria + Nothosauroidea to the exclusion of Pistosauroidea (Li and Liu 2020). Suprisingly, the type specimen of *Hanosaurus* is not recovered in a clade with the referred specimen. Taxonomic distinction of both specimens is supported by the fact that the anteriorly and posteriorly weakly emarginated coracoid of the type specimen of *Hanosaurus* (Rieppel 1998a; pers. obs. of IVPP V 3231) more closely resembles that of *Corosaurus* (Storrs 1991) than the sub-oval coracoid present in the referred specimen of *Hanosaurus* (Wang et al. 2022), which likely does not represent *Hanosaurus*, but is probably closely related to or even referable to ‘*Lariosaurus*’ *sanxiaensis* (Cheng et al. 2016b; Li and Liu 2020). *Corosaurus* is recovered as the earliest-diverging eosauropterygian (result similar to that obtained by Rieppel [1994] and Li and Liu [2020]), in contrast to some other phylogenetic analyses, which recovered it as a pistosauroid (Rieppel 1998b) or the earliest-diverging eusauropterygian (Lin et al. 2021). *Wumengosaurus* is recovered as the sister-taxon of the clade comprising pachypleurosaurs, nothosaurs and pistosaurs, in contrast to a recent phylogenetic analysis which recovered it wtihin pachypleurosaurs (Xu et al. 2022), but in agreement with the phylogenetic results obtained by Wu et al. (2011) that recovered *Wumengosaurus* as the outgroup to a clade comprising pachypleurosaurs and nothosaurs.

### The early evolutionary assembly of the sauropterygian body plan

Our phylogenetic analysis indicates that *Eusaurosphargis* (and *Palatodonta*) likely represent the morphology of the last common terrestrial ancestor of sauropterygians, indicating it possessed well-developed dermal armor and the characteristic pectoral girdle and pes morphology that underwent further modifications in Sauropterygia. The topology recovered by our phylogenetic analysis demonstreates that the early evolution of sauropterygians first involved diversification within a shallow marine environment and exploration of various food resources, as evidenced by the disparate morphologies represented by *Atopodentatus* (herbivore), placodonts (durophages), saurosphargids and early-diverging members of the eosauropterygian lineage (likely feeding on fish and invertebrates). Three key episodes can be identified in the evolution of the eosauropterygian body plan. The first, represented by *Pomolispondylus*, involved a reduction of osteoderms and shortening of the transverse processes of the dorsal neural spines. *Majiashanosaurus* and the referred specimen of *Hanosaurus hupehensis*/’*Lariosaurus*’ *sanxiaensis* represent the second phase, in whch the osteoderms underwent complete reduction. The type specimen of *Hanosaurus hupehensis* represents the earliest stage of the evolution of the characteristic eosauropterygian pectoral girdle morphology with an anteriorly and posteriorly emarginated coracoid. *Corosaurus*, the earliest true eosauropterygian, differs from *Hanosaurus* most notably in the presence of a skull with a preorbital region shorter than the postorbital region, indicating a change in cranial biomechanics at the base of eosauropterygia. The presence of the stratigraphically oldest saurosphargid (*Prosaurosphargis*), stratigraphically oldest representatives of the lineage leading to Eosauropterygia (*Pomolispondylus, Hanosaurus*, ‘*Lariosaurus*’ *sanxiaensis, Majiashanosaurus*), and the earliest-diverging placodontiform *Atopodentatus* in the Early–Middle Triassic of South China (Figure 8) indicates that sauropterygians likely originated and underwent rapid diversification in South China in the aftermath of the end-Smithian extinction, similar to ichthyosauromorphs (Motani et al. 2017; Moon and Stubbs 2020), but well-constrained stratigraphic data for early sauropterygians are needed to further test this hypothesis.

Our phylogenetic analysis indicates the important role of body armor in sauropterygian evolution, which was likely a preadaptation that allowed colonisation of the shallow marine realm by a *Eusaurosphargis*-like ancestor. Further elaboration of the armor in the shallow marine placodonts and saurosphargids, perhaps as response to predation pressure, and the reduction and complete loss of dermal armor in the lineage leading to Eosauropterygia, likely involved with the evolution of an efficient swimming style, demonstrate striking parallels with the evolution of another important group of Mesozoic marine reptiles – the ichthyosaurs. Early-diverging representatives of Ichthyosauromorpha – hupehsuchians and omphalosaurids – also possesed a covering of osteoderms superficially similar to that present in *Eusaurosphargis*, early-diverging placodonts and saurosphargids (Chen et al. 2014b; Motani et al. 2016; Qiao et al. 2022). In the ichthyosauromorph lineage, the osteoderm covering was completely lost in *Chaohusaurus*, a basal ichthyosauriform that evolved an anguilliform mode of swimming and most likely a pelagic lifestyle (Motani et al. 1996; Nakajima et al. 2014; Huang et al. 2019; Qiao et al. 2022). This indicates that the evolutionary reduction of dermal armor in both sauropterygians and ichthyosauromorphs followed a similar, striking pattern, seemingly in response to increasing adaptation to an aquatic lifestyle. These evolutionary parallels seem to demonstrate that osteoderm dermal body armor could have been a possible prerequsite (preadaptation) for the invasion of the shallow marine realm in different diapsid clades. Originally functioning as one of the adaptations to counteract buoyancy in early-diverging marine reptiles, osteoderms were eventually lost during the aquisition of active swimming (Houssaye 2009). Fossils of terrestrial ancestors of ichthyosauromorphs and thalattosaurs are needed to further test this evolutionary scenario.

### Phylogenetic interrelationships within Diapsida

Our phylogenetic analysis is thus far the first phylogenetic analysis based on a morphology-only dataset of Diapsida that recovers a close relationship between Archosauromorpha and Testudinata within a monophyloetic Archelosauria. A close phylogenetic relationship between Archosauromorpha and Testudinata has been established on the basis of molecular data in the last twenty years but was previously not recovered by any phylogenetic analysis based entirely on morphological data, in which turtles were usually recovered as more closely related to lepidosauromorphs than archosauromorphs (reviewed in Lyson and Bever 2020). Furthermore, our analysis recovers *Eunotosaurus* from the Permian of South Africa outside of Sauria, in contrast to all other recent phylogenetic analyses focussing on the phylogenetic interrelationships among Reptilia, which recovered it as an early-diverging stem turtle (Lyson et al. 2013; Bever et al. 2015; Li et al. 2018; Schoch and Sues 2018). This indicates that the characteristic morphological features of the skull and postcranial skeleton shared between *Eunotosaurus* and early turtles, such as elongate vertebrae and broadened ribs, evolved convergently in both taxa. Our results demonstrate the importance of including not only a broad sample of clades in phylogenetic analyses, but also their stratigraphically oldest and/or anatomically most plesiomorphic representatives, in order to accurately reconstruct the phylogenetic relationships between major lineages of extant and extinct tetrapods.

### Conclusions

The new saurosphargid *Prosaurosphargis yingzishanensis* gen. et sp. nov. from the Early Triassic of South China represents the earliest reported occurrence of Saurosphargidae, extending their temporal range back by 4.5 Ma. An updated phylogenetic analysis recovers saurosphargids as nested within Sauropterygia, forming a clade with Eosauropterygia to the exclusion of Placodontia. A clade comprising *Eusaurosphargis* and *Palatodonta* forms the sister-group to Sauropterygia within Sauropterygomorpha tax. nov. and their morphology likely represents the morphology of the last common terrestrial ancestor of Sauropterygia. The herbivorous sauropterygian *Atopodentaus* is recovered within Placodontiformes, whereas *Pomolispondylus, Hanosaurus* and *Majishanosaurus* are recovered in a grade leading to Eosauropterygia, with the type and referred specimens of *Hanosaurus* likely representing distinct taxa. Our new phylogenetic hypothesis indicates sauropterygians originated and diversified in South China in the aftermath of the Permo-Triassic mass extinction event and suggests an important role of dermal armor in the early evolution of the group. Three major marine reptile clades – Sauropterygomorpha, Ichthyosauromorpha and Thalattosauria are recovered in a clade with Archosauromorpha and Testudinata, comprising a monophyletic Archelosauria.

## Materials and Methods

In order to investigate the phylogenetic position of HFUT YZSB-19-109, the specimen was scored into a modified version of a data matrix focusing on the phylogenetic interrelationships between the major groups of diapsid reptiles published by Qiao et al. (2022), which in itself is a modified version of the data matrix previously published by Jiang et al. (2016) and Chen et al. (2014). The data matrix contains 58 OTUs (Operational Taxonomic Units) scored for a total of 221 characters – characters 1–220 are the original characters of Qiao et al. (2022) and character 221 was adapted from Li et al. 2014 (ch. 88) (Supplementary File 1). The following nine OTUs were added to the original 44 OTUs included in the dataset of Qiao et al. (2022) with the aim of including all currently known Early Triassic sauropterygians and increasing the sampling of early-diverging representatives of the main sauropterygian lineages, as well as their potential sister-groups: *Pappochelys rosinae* (scored after Schoch and Sues 2015, 2018); *Odontochelys semitestacea* (Li et al. 2008); *Hanosaurus hupehensis* (type specimen) (Rieppel 1998a and personal observation of specimen IVPP V 3231), *Hanosaurus hupehensis* (referred specimen) (Wang et al. 2022), *Majiashanosaurus discocoracoidis* (Jiang et al. 2014), *Corosaurus alcovensis* (Storrs 1991; Rieppel 1998b), *Atopodentatus unicus* (Cheng et al. 2014; Li et al. 2016), *Palatodonta bleekeri* (Neenan et al. 2013), *Paraplacodus broilii* (Peyer 1935; Rieppel 2000b), *Pomolispondylus biani* (Cheng et al. 2022) and *Prosaurosphargis yingzishanensis* gen. et sp. nov. (HFUT YZSB-19-109).

Parsimony analysis of the data matrix was performed in TNT 1.5 (Goloboff and Catalano 2016) using a Traditional Search algorithm (random seed =1, replications of Wagner trees = 1000), followed by an additional round of TBR branch-swapping. All characters were treated as equally weighted and unordered.

## Acknowledgements

We thank P.M. Sander for early discussions, W. Lin for sharing photographs of the *Hanosaurus* holotype, HFUT Paleontology Lab members for assistance in the field, and the Willi Hennig Society for making the program TNT publically available. F. Huang, L. Li and T. Sato skillfully prepared the fossil specimen and T. Hollman produced the skeletal reconstruction in Fig. 2B. The research was supported by the National Natural Science Foundation of China (42172026 and 41772003 to J.L., 42202006 to A.S.W., and 41902104 to Y.S.), the Fundamental Research Funds for the Central Universities of China (PA2020GDKC0022 to J.L.), the Department of Natural Resources of Anhui Province (2021-g-2-16 to J.L.), the China Postdoc Council (Postdoctoral Fellowship to A.S.W.), and the China Scholarship Council (202106690044 to Y.Q.).

## Data availability

All data generated or analysed during this study are included in the manuscript and supporting files.

## References

Bardet, N. 1994. Extinction events among Mesozoic marine reptiles. Historical Biology, 7(4), 313–324.

Benson, R.B.J. 2013. ‘Marine reptiles’ in MacLeod, N., Archibald, J. D., and Levin, P. (eds) Grzimek’s Animal Life Encyclopedia: Extinction. Gale Research Inc., Farmington Hills, Michigan, pp. 267-279.

Benton, M.J. 2015. When life nearly died: The greatest mass extinction of all time. 2nd edition. Thames & Hudson, London, 352 pp.

Bureau of Geology and Mineral Resources of Hubei Province 1990. Regional Geology of Hubei Province. Geological Publishing House, Beijing. 705 p.

Carroll, R.L. and Gaskill, P. 1985. The nothosaur Pachypleurosaurus and the origin of plesiosaurs. Philosophical Transactions of the Royal Society of London. B, Biological Sciences, 309(1139), 343–393.

Chen, C., Chen, X., Cheng, L., and Yan, C. 2016a. Nangzhang-Yuan’an Fauna, Hubei Province and its significance for biotic recovery. Acta Geologica Sinica, 90, 409-420 (in Chinese with English abstract).

Chen, X., Cheng, L., Wang, C., and Zhang B. 2016b. Triassic Marine Reptile Faunas from Middle and Upper Yangtze Areas and Their Coevolution with Environment. Geological Publishing House, Beijing, 234p (in Chinese with English abstract).

Chen, X.H., Motani, R., Cheng, L., Jiang, D.Y. and Rieppel, O. 2014. The enigmatic marine reptile Nanchangosaurus from the Lower Triassic of Hubei, China and the phylogenetic affinities of Hupehsuchia. PLoS One, 9(7), e102361.

Chen, X., Sander, P.M., Cheng, L., and Wang, X. 2013. A new Triassic primitive ichthyosaur from Yuanan, South China. Acta Geologica Sinica (English Edition), 87, 672-677.

Chen, Y., Jiang, H., Ogg, J.G., Zhang, Y., Gong, Y. and Yan, C., 2020. Early-Middle Triassic boundary interval: Integrated chemo-bio-magneto-stratigraphy of potential GSSPs for the base of the Anisian Stage in South China. Earth and Planetary Science Letters, 530: 115863.

Chen Y.J., Shen, Y.F., and Liu, J. 2022. Carbonate laminites from the Nanzhang-Yuanan fauna in the Lower Triassic Jialingjiang Formation, South China: origins and significances. The 21st International Sedimentological Congress, Beijing.

Cheng, L., Chen, X.H, Zeng, X., and Cai, Y. 2012. A new eosauropterygian (Diapsida: Sauropterygia) from the Middle Triassic of Luoping, Yunnan Province. Journal of Earth Science, 23(1), 33–40.

Cheng, L., Chen, X.H., Shang, Q.H. and Wu, X.C., 2014. A new marine reptile from the Triassic of China, with a highly specialized feeding adaptation. Naturwissenschaften, 101(3), 251–259.

Cheng, L., Yan, C., Chen, X., Zeng, X., and Motani, R. 2015. Characteristics and significance of Nanzhang/Yuanan Fauna, Hubei Province. Geology in China, 42, 676-684 (in Chinese with English abstract).

Cheng, L., C. Moon, B., Yan, C., Motani, R., Jiang, D., An, Z. and Fang, Z., 2022. The oldest record of Saurosphargiformes (Diapsida) from South China could fill an ecological gap in the Early Triassic biotic recovery. PeerJ, 10, e13569.

Crawford, N.G., Parham, J.F., Sellas, A.B., Faircloth, B.C., Glenn, T.C., Papenfuss, T.J., Henderson, J.B., Hansen, M.H. and Simison, W.B. 2015. A phylogenomic analysis of turtles. Molecular phylogenetics and evolution, 83, 250–257.

Crofts, S.B., Neenan, J.M., Scheyer, T.M. and Summers, A.P. 2017. Tooth occlusal morphology in the durophagous marine reptiles, Placodontia (Reptilia: Sauropterygia). Paleobiology, 43(1), 114–128.

de Braga, M. and Rieppel, O. 1997. Reptile phylogeny and the interrelationships of turtles. Zoological Journal of the Linnean Society, 120(3), 281–354.

Drevermann, F.R. 1933. Die Placodontier. 3. Das Skelett von Placodus gigas Agassiz im Senckenberg-Museum. Abhandlungender senckenbergischen naturforschenden Gesellschaft, 38, 319–364.

Goloboff, P.A. and Catalano, S.A. 2016. TNT version 1.5, including a full implementation of phylogenetic morphometrics. Cladistics, 32(3), 221–238.

Hirasawa, T., Nagashima, H. and Kuratani, S. 2013. The endoskeletal origin of the turtle carapace. Nature Communications, 4(1), 1–7.

Hsu, K.J., He, Q., Wu, Y., and Zhu, Z. 1983. Carbon and oxygen stable isotope geochemistry of the carbonates above and below the synchronous marker “Mung Bean” rock between the Early and Middle Triassic in the Southwest China. Journal of Chinese Geological Academy Chengdu Geological Institute, 4, 1-12 (in Chinese with English abstract).

Huang, J.D., Motani, R., Jiang, D.Y., Tintori, A., Rieppel, O., Zhou, M., Ren, X.X. and Zhang, R. 2019. The new ichthyosauriform Chaohusaurus brevifemoralis (Reptilia, Ichthyosauromorpha) from Majiashan, Chaohu, Anhui Province, China. PeerJ, 7, e7561.

Huene, F. v. 1936. Henodus chelyops, ein neuer Placodontier. Palaeontographica A, 84, 99–148.

Jiang, D.Y., Motani, R., Hao, W.C., Rieppel, O., Sun, Y.L., Schmitz, L. and Sun, Z.Y., 2008. First record of Placodontoidea (Reptilia, Sauropterygia, Placodontia) from the eastern Tethys. Journal of Vertebrate Paleontology, 28(3), 904-908.

Jiang, D.Y., Motani, R., Tintori, A., Rieppel, O., Chen, G.B., Huang, J.D., Zhang, R., Sun, Z.Y. and Ji, C., 2014. The early Triassic eosauropterygian Majiashanosaurus discocoracoidis, gen. et sp. nov. (Reptilia, Sauropterygia), from Chaohu, Anhui Province, People’s Republic of China. Journal of Vertebrate Paleontology, 34(5), 1044–1052.

Jiang, D.Y., Motani, R., Huang, J.D., Tintori, A., Hu, Y.C., Rieppel, O., Fraser, N.C., Ji, C., Kelley, N.P., Fu, W.L. and Zhang, R. 2016. A large aberrant stem ichthyosauriform indicating early rise and demise of ichthyosauromorphs in the wake of the end-Permian extinction. Scientific Reports, 6(1), 1–9.

Kelley, N.P., and Pyenson, N.D. 2015. Evolutionary innovation and ecology in marine tetrapods from the Triassic to the Anthropocene. Science, 348(6232), aaa3716.

Kelley, N.P., Motani, R., Jiang, D.Y., Rieppel, O. and Schmitz, L. 2014. Selective extinction of Triassic marine reptiles during long-term sea-level changes illuminated by seawater strontium isotopes. Palaeogeography, Palaeoclimatology, Palaeoecology, 400, 9–16.

Ketchum, H.F., and Benson, R.B.J. 2010. Global interrelationships of Plesiosauria (Reptilia, Sauropterygia) and the pivotal role of taxon sampling in determining the outcome of phylogenetic analyses. Biological Reviews, 85(2), 361–392.

Klein, N., 2009. Skull morphology of Anarosaurus heterodontus (Reptilia: Sauropterygia: Pachypleurosauria) from the Lower Muschelkalk of the Germanic Basin (Winterswijk, The Netherlands). Journal of Vertebrate Paleontology, 29(3), 665–676.

Klein, N., 2012. Postcranial morphology and growth of the pachypleurosaur Anarosaurus heterodontus (Sauropterygia) from the Lower Muschelkalk of Winterswijk, The Netherlands. Paläontologische Zeitschrift, 86(4), 389–408.

Klein, N. and Scheyer, T.M. 2013. A new placodont sauropterygian from the Middle Triassic of the Netherlands. Acta Palaeontologica Polonica, 59(4), 887–902.

Li, C., Rieppel, O., Wu, X.C., Zhao, L.J., and Wang, L.T. 2011. A new Triassic marine reptile from southwestern China. Journal of Vertebrate Paleontology, 31(2), 303–312.

Li, C., Rieppel, O., Cheng, L. and Fraser, N.C. 2016. The earliest herbivorous marine reptile and its remarkable jaw apparatus. Science Advances, 2(5), e1501659.

Li, C., Jiang, D.Y., Cheng, L., Wu, X.C. and Rieppel, O. 2014. A new species of Largocephalosaurus (Diapsida: Saurosphargidae), with implications for the morphological diversity and phylogeny of the group. Geological Magazine, 151(1), 100–120.

Li, C., Fraser, N.C., Rieppel, O. and Wu, X.C. 2018. A Triassic stem turtle with an edentulous beak. Nature, 560(7719), 476–479.

Li, J., Zhao, B., Zou, Y., Chen, G. 2020. Research Status of Hupehsuchia and Preliminary Understanding of Their Environmental Evolution. Resources Environment & Engineering, 34, 485-493 (in Chinese with English abstract).

Li, J., Liu, J., Li, C., and Huang, Z. 2002. The horizon and age of the marine reptiles from Hubei Province, China. Vertebrata Palasiatica, 40, 241-244 (in Chinese with English abstract).

Li, Q. and Liu, J. 2020. An Early Triassic sauropterygian and associated fauna from South China provide insights into Triassic ecosystem health. Communications Biology, 3(1), 1–11.

Lin, W.B., Jiang, D.Y., Rieppel, O., Motani, R., Tintori, A., Sun, Z.Y. and Zhou, M. 2021. Panzhousaurus rotundirostris Jiang et al., 2019 (Diapsida: Sauropterygia) and the recovery of the monophyly of Pachypleurosauridae. Journal of Vertebrate Paleontology, 41(1), e1901730.

Liu, X.Q., Lin, W.B., Rieppel, O., Sun, Z.Y., Li, Z.G., Lu, H., and Jiang, D.Y. 2015. A new specimen of Diandongosaurus acutidentatus (Sauropterygia) from the Middle Triassic of Yunnan, China. Vertebrata Palasiatica, 53(4), 281–290.

Lyson, T.R. and Bever, G.S. 2020. Origin and evolution of the turtle body plan. Annual Review of Ecology, Evolution, and Systematics, 51(1), 143–166.

Lyson, T.R., Bever, G.S., Scheyer, T.M., Hsiang, A.Y. and Gauthier, J.A. 2013. Evolutionary origin of the turtle shell. Current Biology, 23(12), pp.1113–1119.

Martínez, R.N., Simões, T.R., Sobral, G., and Apesteguía, S. 2021. A Triassic stem lepidosaur illuminates the origin of lizard-like reptiles. Nature, 597(7875), 235–238.

Motani, R., 2009. The evolution of marine reptiles. Evolution: Education and Outreach, 2(2), 224–235.

Motani, R., You, H., and McGowan, C. 1996. Eel-like swimming in the earliest ichthyosaurs. Nature, 382(6589), 347–348.

Motani, R. and Vermeij, G.J. 2021. Ecophysiological steps of marine adaptation in extant and extinct non-avian tetrapods. Biological Reviews, 96(5), 1769–1798.

Motani, R., Jiang, D.Y., Chen, G.B., Tintori, A., Rieppel, O., Ji, C. and Huang, J.D., 2015. A basal ichthyosauriform with a short snout from the Lower Triassic of China. Nature, 517(7535), 485–488.

Moon, B.C., and Stubbs, T.L. 2020. Early high rates and disparity in the evolution of ichthyosaurs. Communications Biology, 3(1), 1–8.

Neenan, J.M., Klein, N. and Scheyer, T.M. 2013. European origin of placodont marine reptiles and the evolution of crushing dentition in Placodontia. Nature Communications, 4(1), 1–8.

Neenan, J.M., Li, C., Rieppel, O. and Scheyer, T.M. 2015. The cranial anatomy of Chinese placodonts and the phylogeny of Placodontia (Diapsida: Sauropterygia). Zoological Journal of the Linnean Society, 175(2), 415–428.

Nosotti, S. and Rieppel, O. 2003. Eusaurosphargis dalsassoi n. gen n. sp., a new, unusual diapsid reptile from the Middle Triassic of Besano (Lombardy, N Italy). Memorie della Società Italiana di Scienze Naturali e del Museo Civico di Storia Naturale di Milano, 31, 3–33.

Owen, R. 1860. Palaeontology; or, a systematic summary of extinct animals and their geologic remains. Adam and Charles Black, Edinburgh.

Peyer, B. 1935. Die Triasfauna der Tessiner Kalkalpen. VIII. Weitere Placodontierfunde. Abhandlungen der schweizerischen Palaontologischen Gesellschaft, 55, 1–26.

Qiao, Y. Liu, J., Wolniewicz, A.S., Iijima, M., Shen, Y., Wintrich, T., Li, Q. and Sander, P.M. 2022. A globally distributed durophagous marine reptile clade supports the rapid recovery of pelagic ecosystems after the Permo-Triassic mass extinction. Communications Biology, in press.

Rieppel, O. 1989a. Helveticosaurus zollingeri Peyer (Reptilia, Diapsida) skeletal paedomorphosis, functional anatomy and systematic affinities. Palaeontographica. Abteilung A. Paläozoologie, Stratigraphie, 208(4-6), 123-152.

Rieppel, O. 1989b. A new pachypleurosaur (Reptilia: Sauropterygia) from the Middle Triassic of Monte San Giorgio, Switzerland. Philosophical Transactions of the Royal Society of London. B, Biological Sciences, 323(1212), 1–73.

Rieppel, O. 1994. Osteology of Simosaurus gaillardoti, and the phylogenetic interrelationships of stem-group Sauropterygia. Fieldiana (Geology), n.s. 28, 1–85.

Rieppel, O. 1995. The genus Placodus: Systematics, Morphology, Paleobiogeography, and Paleobiology. Fieldiana (Geology), n.s. 31, 1–44.

Rieppel, O. 1998a. The systematic status of Hanosaurus hupehensis (Reptilia, Sauropterygia) from the Triassic of China. Journal of Vertebrate Paleontology, 18(3), 545–557.

Rieppel, O. 1998b. Corosaurus alcovensis Case and the phylogenetic interrelationships of Triassic stem-group Sauropterygia. Zoological Journal of the Linnean Society, 124(1), 1–41.

Rieppel, O. 2000a. ‘Sauropterygia I’ in Wellnhofer P. (ed) Encyclopedia of Paleoherpetology, Volume 12A, Verlag Dr. Friedrich Pfeil, Munich, pp. 1-134.

Rieppel, O. 2000b. Paraplacodus and the phylogeny of the Placodontia (Reptilia: Sauropterygia). Zoological Journal of the Linnean Society, 130(4), 635–659.

Rieppel, O. 2002. Feeding mechanics in Triassic stem-group sauropterygians: the anatomy of a successful invasion of Mesozoic seas. Zoological Journal of the Linnean Society, 135(1), 33–63.

Rieppel, O., and Lin, K. 1995. Pachypleurosaurs (Reptilia: Sauropterygia) from the Lower Muschelkalk, and a review of the Pachypleurosauroidea. Fieldiana (Geology), n.s. 32, 1–44.

Sander, P.M., Rieppel, O.C. and Bucher, H. 1997. A new pistosaurid (Reptilia: Sauropterygia) from the Middle Triassic of Nevada and its implications for the origin of the plesiosaurs. Journal of Vertebrate Paleontology, 17(3), 526–533.

Sander, P.M., Griebeler, E.M., Klein, N., Juarbe, J.V., Wintrich, T., Revell, L.J. and Schmitz, L. 2021. Early giant reveals faster evolution of large body size in ichthyosaurs than in cetaceans. Science, 374(6575), eabf5787.

Scheyer, T.M. 2007. Skeletal histology of the dermal armor of Placodontia: the occurrence of ‘postcranial fibro-cartilaginous bone’ and its developmental implications. Journal of Anatomy, 211(6), 737–753.

Scheyer, T.M., Romano, C., Jenks, J., and Bucher, H. 2014. Early Triassic marine biotic recovery: the predators’ perspective. PLoS One, 9(3), e88987.

Scheyer, T.M., Klein, N., Sichelschmidt, O., Neenan, J.M. and Albers, P., 2019. With plates and spikes-the heavily armoured Eusaurosphargis aff. dalsassoi. Staringia, 16(5/6), 216–222.

Scheyer, T.M., Neenan, J.M., Bodogan, T., Furrer, H., Obrist, C., and Plamondon, M. 2017. A new, exceptionally preserved juvenile specimen of Eusaurosphargis dalsassoi (Diapsida) and implications for Mesozoic marine diapsid phylogeny. Scientific Reports, 7(1), 1–22.

Scheyer, T.M., Oberli, U., Klein, N. and Furrer, H. 2022. A large osteoderm-bearing rib from the Upper Triassic Kössen Formation (Norian/Rhaetian) of eastern Switzerland. Swiss Journal of Palaeontology, 141(1), 1–9.

Schoch, R.R., and Sues, H.D. 2015. A Middle Triassic stem-turtle and the evolution of the turtle body plan. Nature, 523(7562), 584–587.

Schoch, R.R. and Sues, H.D. 2018. Osteology of the Middle Triassic stem-turtle Pappochelys rosinae and the early evolution of the turtle skeleton. Journal of Systematic Palaeontology, 16(11), 927–965.

Shang, X.H., Wu, X.C., and Li, C. 2011. A new eosauropterygian from Middle Triassic of eastern Yunnan Province, southwestern China. Vertebrata PalAsiatica, 49(2), 155–171.

Simoes, T.R., Caldwell, M.W., Tałanda, M., Bernardi, M., Palci, A., Vernygora, O., Bernardini, F., Mancini, L. and Nydam, R.L. 2018. The origin of squamates revealed by a Middle Triassic lizard from the Italian Alps. Nature, 557(7707), 706–709.

Storrs, G.W. 1991. Anatomy and relationships of Corosaurus alcovensis (Diapsida: Sauropterygia) and the Triassic Alcova limestone of Wyoming. Bulletin of the Peabody Museum of Natural History, 44, 1–151.

Stubbs, T.L., and Benton, M.J. 2016. Ecomorphological diversifications of Mesozoic marine reptiles: the roles of ecological opportunity and extinction. Paleobiology, 42(4), 547–573.

Tschanz, K. 1989. Lariosaurus buzzii n. sp. from the Middle Triassic of Monte San Giorgio (Switzerland) with comments on the classification of nothosaurs. Palaeontographica. Abteilung A. Paläozoologie, Stratigraphie, 208(4-6), 153–179.

Vermeij, G.J. and Motani, R. 2018. Land to sea transitions in vertebrates: the dynamics of colonization. Paleobiology, 44(2), 237–250.

Wang, J., Cheng, L., and Zeng, X. 2011. Sedimentary characteristics of the second member of Jialingjiang formation in the Lower Triassic of the Middle Yangtze Area and its reflection on the burial environment of marine reptiles. Abstract Volume of the 26^th^ Annual Academic Meeting of the Palaeontological Society of China, Guanling, 159 (in Chinese with English abstract).

Wang, W., Li, C. and Wu, X.C. 2019. An adult specimen of Sinocyamodus xinpuensis (Sauropterygia: Placodontia) from Guanling, Guizhou, China. Zoological Journal of the Linnean Society, 185(3), 910–924.

Wang, W., Ma, F. and Li, C., 2020. First subadult specimen of Psephochelys polyosteoderma (Sauropterygia, Placodontia) implies turtle-like fusion pattern of the carapace. Papers in Palaeontology, 6(2), 251–264.

Wang, W., Shang, Q.H., Cheng, L., Wu, X.C., and Li, C. 2022. Ancestral Body Plan and Adaptive Radiation of Sauropterygian Marine Reptiles. bioRxiv, doi: https://doi.org/10.1101/2022.04.25.489368.

Willemse, D.M., Willemse, N.W., Voeten, D.F.A.E. 2019. An aff. Eusaurosphargis vertebra associated with an isolated dentary. Staringia, 16(5-6), 273–277.

Wu, X.C., Cheng, Y.N., Li, C., Zhao, L.J. and Sato, T. 2011. New information on Wumengosaurus delicatomandibularis Jiang et al., 2008 (Diapsida: Sauropterygia), with a revision of the osteology and phylogeny of the taxon. Journal of Vertebrate Paleontology, 31(1), 70–83.

Xu, G.H., Ren, Y., Zhao, L.J., Liao, J.L. and Feng, D.H. 2022. A long-tailed marine reptile from China provides new insights into the Middle Triassic pachypleurosaur radiation. Scientific Reports, 12(1), 1–12.

Yan, C., Cheng, L., Tintori, A., Li, Z., and Yang, B. 2018. Stratigraphic characterization and paleoenvironmental analysis of Early Triassic Nanzhang-Yuan’an Fauna, western Hubei Province. Abstract Volume of Joint Meetings on the 12^th^ National Congress and 29^th^ Annual Academic Meeting of the Palaeontological Society of China, Jiaozuo, 237-238 (in Chinese with English abstract).

Yan, C., Li, J., Cheng, L., Zhao, B., Zou, Y., Niu, D., Chen, G., and Fang, Z. 2021. Strata characteristics of the Early Triassic Nanzhang-Yuan’an Fauna in Western Hubei Province. Earth Science, 46, 122-135 (in Chinese with English abstract).

Young C.C. 1965. On the new nothosaurs from Hupeh and Kweichou, China. Vertebrata PalAsiatica, 9, 337–356.

Young, C.C. 1972. A marine lizard from Nanchang, Hupeh Province. Memoirs of the Institute of Vertebrate Paleontology and Paleoanthropology, Academia Sinica, 9, 17–28.

Zhao, B., Zhou, Y., Chen, G., Li, J., Wu, K., Wan, S., and Xu, X. 2022. New research progress on a specimen of Chaohusaurus zhangjiawanensis (diapsida: ichthyosauromorpha). Acta Geologica Sinica (100^th^ Anniversary), 96, 769–782 (in Chinese with English abstract).

Zhao, X., Tong, J., Yao, H., Zhang, K., and Chen, Z. 2008. Anachronistic facies in the Lower Triassic of South China and their implications to the ecosystems during the recovery time. Science in China Series D: Earth Sciences, 51, 1646-1657. doi: 10.1007/s11430-008-0128-y.

